# Elevated *Plasmodium falciparum* sexual conversion in HbAC and HbAS red blood cells in naturally infected malaria patients

**DOI:** 10.1101/2025.11.15.688532

**Authors:** Yasmina Drissi-El Boukili, Vera Kühne, H. Magloire Natama, Pieter Moris, Elisabet Tintó-Font, Odin Goovaerts, Pieter Guetens, Ana Moreno-Murillo, Yarno Valgaerts, T. Edwig Traoré, D. Florence Ouedraogo, Halidou Tinto, Ann Van de Velde, Alfred Cortés, Guy Caljon, Anna Rosanas-Urgell

## Abstract

Malaria transmission relies on the differentiation of asexual parasites into gametocytes, a process initiated by sexual conversion (SC). Mutations in the host hemoglobin beta (HBB) gene are known to influence parasite growth and disease outcome, yet their impact on SC remains unclear. We investigated the effect of HBB mutant genotypes on *Plasmodium falciparum* SC and humoral immunity in individuals from Nanoro, Burkina Faso. To measure SC rates in natural human malaria infections, we developed a new *ex vivo* SC assay (*ev*SCA). We found that in human natural *P. falciparum* infections, SC rates were higher in individuals with HbAS or HbAC than in individuals with wild-type HBB (HbAA). Consistently, using an *in vitro* SC assay (*iv*SCA) based on the NF54-*gexp02-Tom* reporter line we found that cultures grown in HbAS red blood cells (RBCs) had higher SC rates than those grown in HbAA RBCs. Furthermore, IgG and IgM responses against trophozoite- and stage I gametocyte-infected RBC antigens, quantified by flow cytometry, did not differ between plasma from individuals with different HBB genotypes. These results demonstrate that exposure to RBCs with HBB mutations enhances SC, highlighting a host-genetic factor that may influence malaria transmission potential.

**AUTHOR SUMMARY:** Transmission of *Plasmodium falciparum* –a malaria causing parasite− depends on the parasite’s ability to produce gametocytes, the stage that infects mosquitoes. In many African regions where malaria is common, hemoglobin (Hb) variants such as hemoglobin S (the sickle-cell trait) and C are highly prevalent reflecting their role in protecting against severe disease. However, it remains unclear whether these variants also influence the parasite’s early shift toward gametocyte development (*i.e.,* sexual conversion [SC]). In this study, we combined *ex vivo* and *in vitro* SC assays to measure SC in red blood cells with different hemoglobin genotypes. We found that SC was higher in parasites growing in red blood cells carrying hemoglobin AC and AS genotypes compared to normal AA hemoglobin. These findings provide experimental support for earlier epidemiological observations reporting that individuals with these hemoglobin variants often carry more gametocytes and are more infectious to mosquitoes. We also measured antibody levels in individuals with mutant RBCs and found no differences in immunity that could explain the variation in SC. Our results show that host genetic background can influence gametocyte development and may shape the infectious reservoir in high-burden settings. This knowledge is important for designing effective strategies for malaria elimination.

## INTRODUCTION

*Plasmodium falciparum* (*P. falciparum*) causes over half a million deaths annually, mostly in young children from low- and middle-income countries. Despite the substantial progress in reducing the global malaria burden during the last decade, this decline has stalled, with some countries reporting a resurgence in clinical cases [1]. Innovative approaches for malaria control and elimination are thus urgently needed.

*Plasmodium* infections have exerted strong evolutionary pressure on the human genome, with hemoglobin mutations providing a well-documented example [2–5]. Mutations in the human HBB gene, such as hemoglobin S (HbS) and hemoglobin C (HbC), result from a substitution of glutamic acid at position 6 with valine in HbS and lysine in HbC. Individuals can carry homozygous (HbSS or HbCC), heterozygous (HbAS or HbAC) or compound heterozygous (HbSC) mutations [6–8]. HBB mutations lead to significant changes in RBC morphology and function, influencing malaria susceptibility and clinical outcomes [5, 9, 10]. HbS mutations induce hemoglobin polymerization under low oxygen, causing RBCs to reshape into sickle forms. HbSS causes ∼60-80% RBC sickling [2] with severe complications including vaso-occlusion, hemolysis, and chronic anemia [11, 12]. This results in early-life mortality, with estimates ranging from 50-90% among children in low-and middle income countries [13]. On the other hand, HbAS carriers are typically asymptomatic [2], with fewer sickled RBCs (∼40%) [12, 14]. HbC mutations cause hemoglobin crystallization without sickling but increase blood viscosity and shorten RBC life span [15]. HbCC and HbAC are usually a mild and asymptomatic disorder, with HbCC occasionally causing mild chronic hemolytic anemia in some individuals [2, 7, 15]. HbSC carriers display moderate disease severity and a mixed phenotype with ∼50% sickled/crystallized and “pita-shaped” RBCs [2, 11, 16].

HbSS is relatively rare (∼2%), likely due to its high mortality, whereas HbCC, although a mild disorder, is also relatively uncommon (∼2%). HbAS and HbAC can reach frequencies of 10-30% in regions such as Mali and Burkina Faso, which has been linked to the protection they confer against *P. falciparum* malaria [4, 5, 8, 17–20]. HbAC offers mild protection against severe malaria, whereas HbAS and HbCC provide strong protection against both uncomplicated and severe malaria [5, 15, 21–23]. How different HBB mutations protect against malaria is not fully understood, but several mechanisms have been proposed [4, 24–31]. HbS infected-RBCs (iRBCs), especially in HbSS, exhibit elevated heme and HO-1−mediated oxidative stress. This causes polymerization, disrupts the actin cytoskeleton, alters iRBCs morphology, and impairs the surface expression of *P. falciparum* erythrocyte membrane protein 1 (PfEMP1) on iRBCs [32–34]. In HbAS iRBCs, this reduces cytoadherence and sequestration, promoting splenic clearance and controlled parasite infection [25, 35].

Beyond protection from disease, HbS and HbC RBCs may influence the transmission potential of *P. falciparum*. Population-based studies have reported conflicting evidence regarding whether HbAS influences gametocyte carriage. Some studies reported no significant association between HbAS and increased gametocyte carriage, while HbAC and HbCC were associated with increased gametocyte carriage in the same [36] and other populations [37]. In contrast, other studies found a link between HbAS and gametocyte carriage [38–40]. Some studies have also reported higher infectivity to mosquitoes in individuals carrying HbAS [36, 37, 40], HbAC [36, 37] and HbCC [36] compared to wild-type HbAA iRBCs.

Gametocyte production is initiated by a process called sexual conversion (SC), in which a variable but typically small proportion (∼1%) of asexual *P. falciparum* parasites commit to sexual development at each replication cycle, a proportion referred to as the SC rate (SCR), and initiate gametocytogenesis [41–43]. Sexual commitment is marked by epigenetic activation of the master regulator *pfap2g*, a transcription factor that irreversibly results in SC and gametocyte development [44–47]. Sexual rings circulate in blood [48–51], while maturation of gametocyte stages I to IV occurs mainly in the bone marrow over ∼10 days [52]. Mature male and female stage V gametocytes re-enter the peripheral blood, resulting in gametocyte carriage, and are the only parasite stages capable of infecting *Anopheles* mosquitoes. Gametocyte carriage determines human to mosquito transmission and correlates with infectivity to mosquitoes [53–58].

The elevated gametocyte carriage reported in individuals with HbAC [36, 37] and, in some studies, HbAS genotypes [38–40] could result from elevated SC but also from prolonged infections, leading to accumulation of gametocytes in peripheral blood over numerous replication cycles. Several host related factors have been shown to increase SC [59], including lysophosphatidylcholine depletion [60–62], reticulocyte-rich environments [63, 64], and antimalarial treatment [65–68]. However, other studies did not find enhanced SC after antimalarial treatment [67]. To date, the link between SC and HBB genetics has been explored to a limited extent, with one study reporting that increased intracellular heme in HbS iRBCs promotes SC [32]. Another potential driver of elevated SC in HBB mutant RBCs is the host immunity against *P. falciparum*. HbS and HbC mutations appear to modulate host immunity against *P. falciparum* by increasing regulatory cytokines (*e.g.,* IL-10) [30], upregulating HO-1 [28], and dampening pro-inflammatory responses (*e.g.,* TNF-α, IFN-γ) [24], which both protect against severe malaria and enhance parasite clearance. While there is evidence of a differential humoral immune response between different HBB genotypes, differences were only detectable in certain populations and for very specific antigens [69–73].

Discrepancies in gametocyte carriage studies highlight the need to investigate upstream processes such as SC. Determining if HBB mutant RBCs enhance *P. falciparum* SC could confirm that carriers of these mutant RBCs are key contributors to the infectious reservoir – as proposed by mosquito feeding studies [36, 37, 40], informing strategies to reduce transmission.

In this study, we investigated the impact of host HBB mutant genotypes (HbAA, HbAC, HbAS, HbSS, HbCC and HbSC) on SC in naturally infected malaria patients and *in vitro P. falciparum* cultures and on IgG and IgM reactivity against trophozoite- and stage I gametocyte-infected RBCs (iRBCs) by flow cytometry.

## RESULTS

### HbAC and HbAS genotype prevalences in Nanoro, Burkina Faso

Using Sanger Sequencing, the HBB gene was genotyped in participants of a longitudinal cohort study (*i.e., InHost* study [74]) in Nanoro, Burkina Faso (*n* = 864) and the population frequencies of HbAA, HbAC, HbCC, HbAS, HbSS, and HbSC genotypes were determined (Fig 1). HbAA (wild-type) was the most frequent (71.5%), followed by HbAC (20%), HbAS (7.7%), HbSC (0.8%), and HbCC (0.2%). No HbSS individuals were identified.

**Fig 1.**
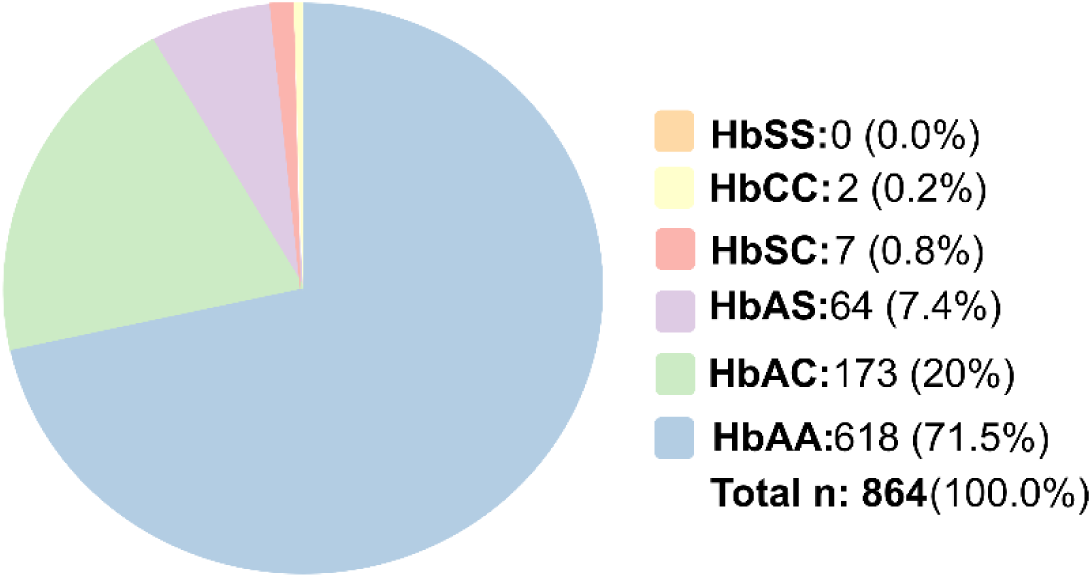
Genotype frequencies of the HBB gene in the study population. Pie chart depicting the total number of individuals (n) and frequencies (%) of the different HBB genotypes in the study population.

### Novel *ex vivo* assay reveals elevated sexual conversion in HbAS/HbAC

To investigate the effect of HBB genotypes on *P. falciparum* SC we measured SCRs *in vivo* in *P. falciparum* isolates collected from naturally infected individuals with different HBB genotypes (14 HbAA, 12 HbAC, and 6 HbAS) living in the malaria-endemic area of Nanoro, Burkina Faso (Fig 2). For this, we developed a new *ex vivo* SC assay (*ev*SCA), optimized and validated as described in *S1 Appendix*. In brief, to assess the proportion of circulating rings in infected individuals that are sexual or asexual, parasites were cultured *ex vivo* in the presence of the PKG inhibitor ML10 (to prevent reinvasion [75]) until the majority of parasites reached the schizont stage. At this time, sexual rings have progressed to Pfs16-positive stage I gametocytes. Next, parasites were concentrated using magnetic columns, and thin blood smears were prepared from the enriched fractions. SCRs were quantified by immunofluorescent staining (IFAs) detecting the proportion of parasites that stained positive for Pfs16 (stage I gametocytes) [76–78] relative to the total number of parasites, stained for DNA/parasite nuclei (the total parasite population consisting of stage I gametocytes plus schizont stages) [45, 79, 80]. The experimental set-up of the *ev*SCAs and representative IFA images are shown in Figs 2A and 2B, respectively. The median SCR was 2.65% (IQR 1.72) in *P. falciparum* isolates from HbAA genotype, 5.05% (IQR 2.55) for HbAC and 18.40% (IQR 19.70) for HbAS genotypes. SCR increased modestly in HbAC (beta-binomial generalized linear mixed model [GLMM], odds ratio (OR)[95%CI] = 1.82 [1.17−2.82], *p (Bonferroni)* = 0.0032) and strongly in HbAS (beta-binomial GLMM, OR [95%CI] = 8.16 [4.34−15.37], *p (Bonferroni)* < 0.0001) genotypes compared to HbAA (Fig 2C, S1A Table). The fold-change between median SCR of *P. falciparum* isolates in HbAC and HbAS relative to the median SCR of wild-type HbAA RBCs was 2.04 (IQR 0.85) in HbAC and 7.07 (IQR 7.32) in HbAS, reflecting the increased SCRs in HbAC and HbAS RBCs compared to HbAA.

**Fig 2.**
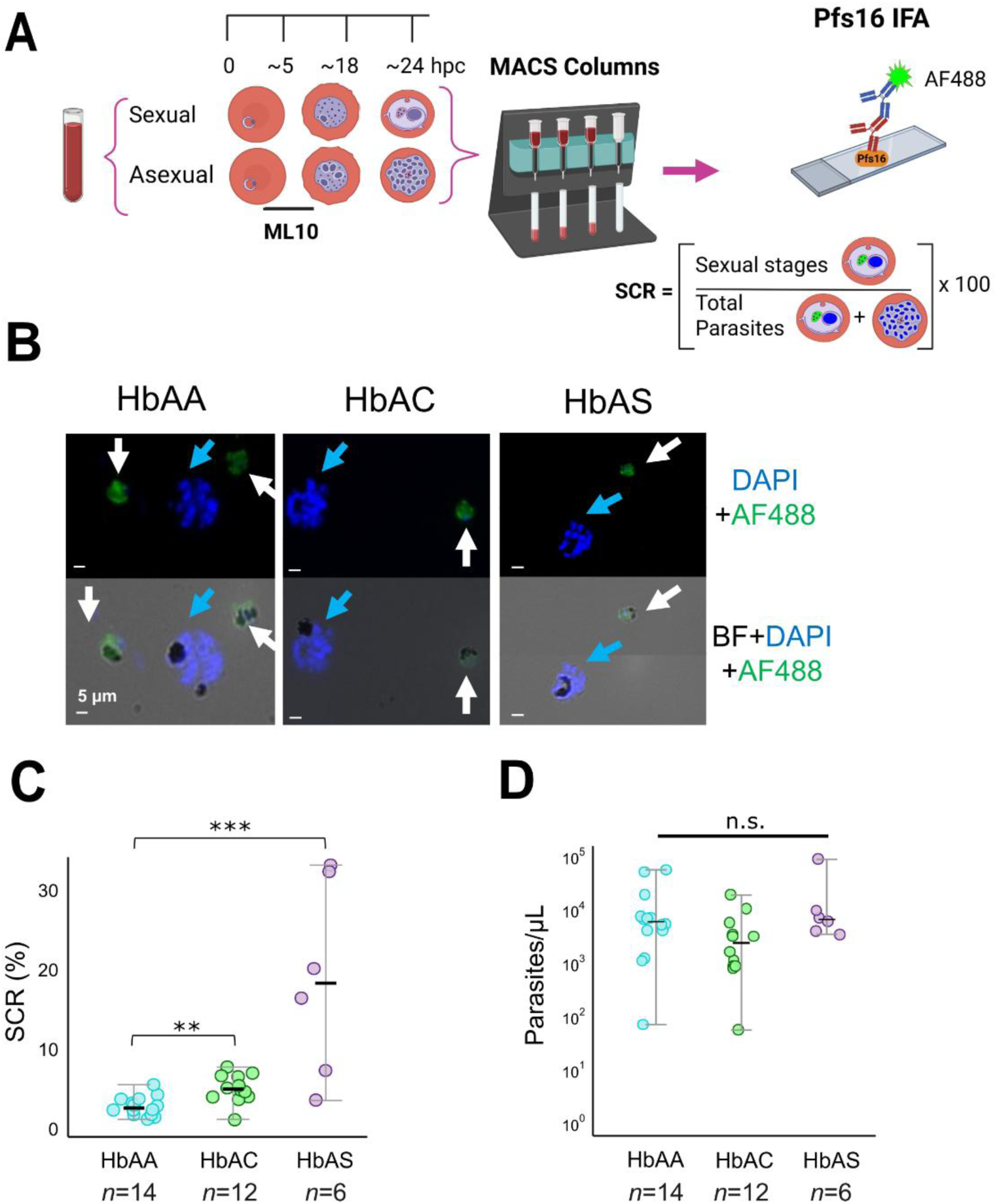
Effect of HbAA, HbAC and HbAS on SC determined with *ex vivo* SC assays. **(A)** Schematic representation of the *ex vivo* SC assays (*ev*SCAs). Parasite isolates from RBCs with different HBB genotypes cultured *ex vivo*, ML10 added at 11 hpi., MACS-enriched stage I gametocytes and schizonts detected by IFA Pfs16 (sexual stages) and DAPI (total parasites) staining. SCR (%): number of sexual stages/number of total parasites. **(B)** Representative IFA images of parasites in HbAA, HbAC and HbAS RBCs. Top row with DAPI (all parasites; blue arrows: schizonts) and AF488 (white arrows: stage I gametocytes) overlay and bottom row including brightfield (BF) channel to visualize the parasite’s hemozoin. Objective: 40×; Scale bar: 5µm. **(C)** SCRs by HBB genotype determined using IFA. Individual dots, represent the mean SCR of three independent technical IFA replicates from one culture; the horizontal line and error bars indicate the median SCR and IQR of all cultures from the same HBB genotype. **(D)** Initial parasitemia at 0 hpc by HBB genotype **Statistical tests: C,** Beta-binomial GL(M)M with per-genotype dispersion parameter and Bonferroni correction for pairwise comparisons; **D,** Kruskal-Wallis test. Significant (*p* < 0.05*, *p* < 0.01**, *p* < 0.001***) and non-significant p-values (ns). **Abbreviations**: AF, Alexa Fluor; DAPI, 4’,6-diamidino-2-phenylindole; GL(M)Ms, Generalized linear (mixed) models; Hb, hemoglobin; HBB, human hemoglobin beta; Hpc, hours post culture; hpi, hours post-invasion; IFA, immunofluorescence assays; ML10, cGMP-dependent protein kinase inhibitor; RBCs, red blood cells; SC, sexual conversion; SCR, sexual conversion rate.

Parasitemias did not significantly differ between mutant HBB RBCs (HbAC and HbAS) and wild-type HbAA RBCs (Kruskal-Wallis test, overall *p* = 0.72, Fig 2D); and SCRs and initial parasitemia did not correlate (Spearman correlation, ρ = 0.05, *p* = 0.62; (S1A Fig). These results indicate that differences in initial parasitemias do not have an impact on the observed elevated SCRs. A risk analysis of different variables in the study population (*i.e.,* gender, village, ethnicity, age, parasitemias, etc.) across HBB genotype groups, showed that SCRs is the only variable that significantly differs amongst the HBB genotypes HbAA, HbAC and HbAS (Kruskal-Wallis test, overall *p* < 0.001; S2 Table).

### *In vitro* sexual conversion in red blood cells of different HBB genotypes

To confirm the findings from the *ev*SCAs, we adapted a previously described *in vitro* SC assay (*iv*SCA) [65] to investigate SC in mutant RBCs (Fig 3A). The assay is based on the *P. falciparum* transgenic reporter line NF54-*gexp02-Tom* [65, 81], which carries an integrated tandem Tomato (tdTom) fluorescent marker under the control of the promoter of the early gametocyte marker gene *gexp02*. Percoll-purified schizonts were grown in cryopreserved RBCs with different HBB genotypes from 28 wild-type and mutant donors, including 15 donors from the malaria-endemic area of Nanoro, Burkina Faso (5 HbAA, 5 HbAC, 4 HbAS, and 1 HbSC) and 13 patients with hemoglobinopathies (12 HbSS and 1 HbSC) from a non-endemic area (Antwerp University Hospital). In addition, non-cryopreserved (fresh) RBCs from 6 control HbAA donors (HbAA CTRL) from a non-endemic area were included (Fig 3A). After 5 h, sorbitol lysis was performed to achieve tight synchronization (5 h age window) and cultures were maintained either with (+Choline) or without (-Choline) a choline supplement, to measure basal or induced SCRs, respectively [60, 79, 81]. At the next cycle (Generation 1),

**Fig 3.**
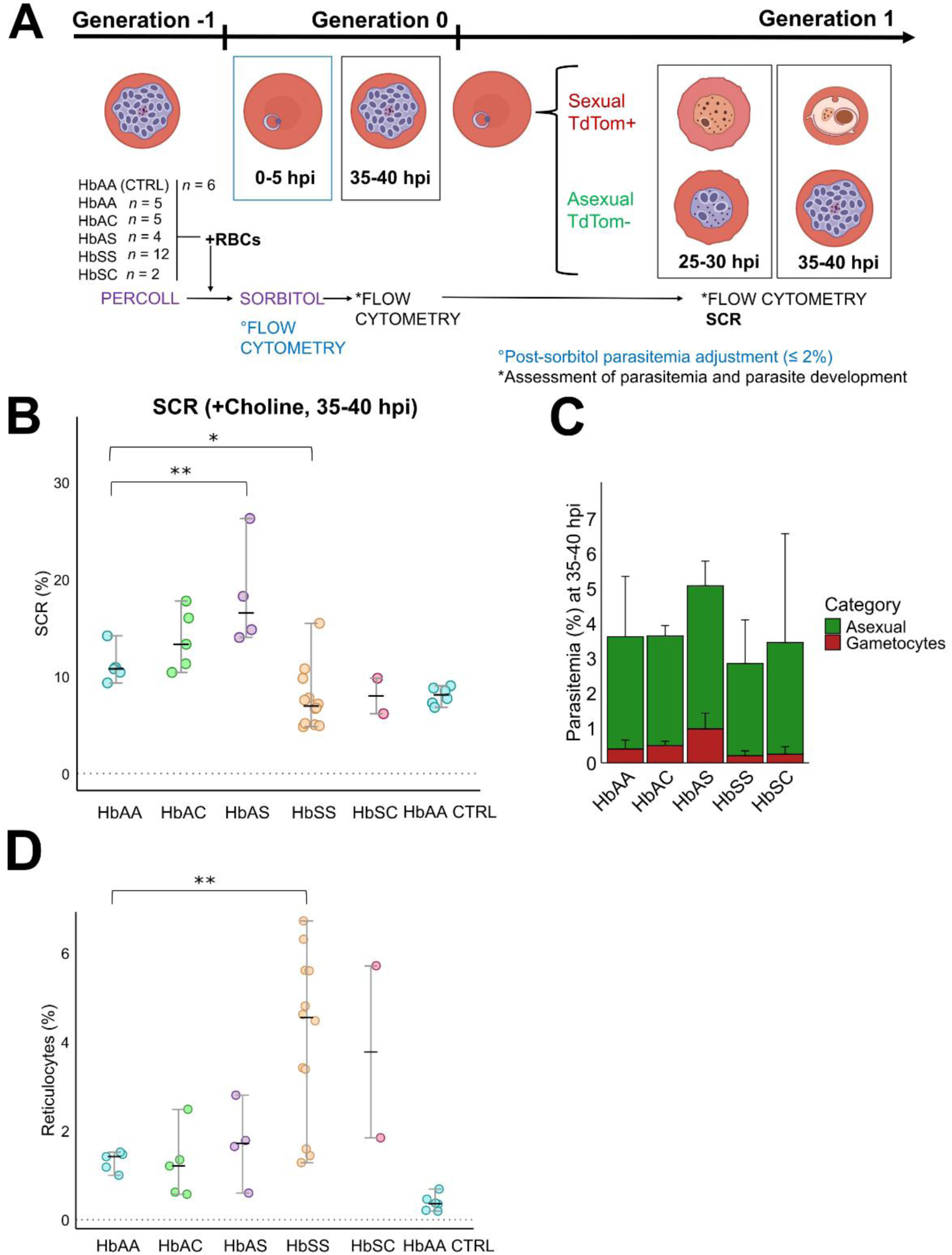
Effect of HbAA, HbAC, HbAS, HbSS and HbSC on SC by *in vitro* SC assays. **(A)** Experimental design of *in vitro* SC assays (*iv*SCAs) using the NF54*-gexp02-Tom P. falciparum* line. Tightly synchronized schizonts (Generation −1) were allowed to reinvade RBCs of different HBB genotypes, and ring stages (0-5 hpi) were cultured under non-inducing (+Choline) and inducing (-Choline) conditions. Parasite development was assessed by flow cytometry at 35-40 hpi (Generation 0) and 25-30 hpi/35-40 hpi (Generation 1). SCRs were measured by flow cytometry at 25-30 hpi and 35-40 hpi (Generation 1). **(B)** SCRs by HBB genotype at 35-40 hpi (Generation 1) in +Choline cultures. **(C)** Proportion of asexual and gametocyte stages and total parasitemia by HBB genotype at 35-40 hpi (Generation 1) in +Choline cultures. **(D)** Reticulocytes percentage by HBB genotype. **Statistical tests:** B and D, Beta-binomial GLMs with Bonferroni correction for pairwise comparisons. Significant *p*-values are shown as: *p* < 0.05*, *p* < 0.01**, *p* < 0.001***; non-significant *p*-values as: ns. **Abbreviations:** CTRL, control sample; GLMs, Generalized linear models; Hb, hemoglobin; HBB, human hemoglobin beta; Hpi, hours post-invasion; SC, sexual conversion; SCR, sexual conversion rates.

SCRs were estimated as the proportion of total parasites (identified by Hoechst33342 staining) that developed as sexual forms (TdTom positive), determined by flow cytometry [81]. A previous report described that the vast majority of sexual parasites expressed the TdTom reporter from 10-15 hpi of Generation 1 onwards [81], and other studies using this assay typically measured SCRs at 20-25 hpi [65]. However, to account for the possible developmental delay observed in cultures grown in RBCs with mutant HBB (see next section), we also measured SCRs at 35-40 hpi. The *iv*SCA assay (Fig 3A, S2 Fig) was validated for use with cryopreserved mutant RBCs as described in the *S1 Appendix*. As detailed in the Methods, analyses were intentionally restricted to parsimonious models due to sample size constraints, and results should be interpreted accordingly.

In non-induced cultures (+Choline), SCRs measured at 35-40 hpi were significantly higher in HbAS RBCs than in wild-type HbAA RBCs (median 16.50% vs. 10.80%, OR[95% CI] = 1.73 [1.12−2.66], *p* [*Bonferroni*] = 0.0056 using beta-binomial GLM) (Fig 3B, S1B and S3 Tables). The same trend was observed for HbAC RBCs, but in this case the difference was not significant (median 13.30%, *p[Bonferroni]* = 0.77 using beta-binomial GLM; S1B Table). In contrast, SCRs were significantly lower in HbSS RBCs (median 6.96%, OR[95% CI] = 0.64 [0.43−0.95], *p* [*Bonferroni*] = 0.021 compared to HbAA RBCs, using beta-binomial GLM) and non-significantly lower in HbSC RBCs (median 7.99%, *p[Bonferroni]* = 0.62, using beta-binomial GLM) (Fig 3B, S1B and S3 Tables). The fold-changes between median SCR in HbAC, HbAS, HbSS and HbSC relative to the median SCR of wild-type HbAA RBCs were 1.24 (IQR 0.44) in HbAC, 1.53 (IQR 0.53) in HbAS, 0.65 (IQR 0.29) in HbSS and 0.74 (IQR 0.17) in HbSC RBCs, reflecting the increased SCRs in HbAC and HbAS RBCs compared to HbAA. The same trends were observed when SCRs were measured at 25-30 hpi, but in this case none of the differences were statistically significant (S3A Fig, S1B and S3 Tables). However, SCRs estimated at 25-30 hpi were generally lower than at 35-40 hpi (S3 Table), supporting the idea that some sexual parasites had not started expressing TdTom yet at 25-30 hpi. Therefore, we considered that, in our experiments with cryopreserved RBCs, measuring SCRs at 35-40 hpi was more appropriate. This is also supported by the results presented in the next section, showing a developmental delay associated with some HBB genotypes.

In cultures in which SC had been stimulated by choline depletion (-Choline), SCRs were significantly lower in cultures with HbSS or HbSC RBCs than in cultures in HbAA RBCs at both 25-30 hpi and 35-40 hpi (beta-binomial GLM; S1B Table). At 25-30 hpi, median SCRs were 24.00% in HbAA cultures, compared with 16.00% in HbSS (OR[95% CI] = 0.52 [0.35−0.79], *p[Bonferroni]* = 0.0003) and 10.20% in HbSC (OR[95% CI] = 0.36 [0.17−0.77], *p[Bonferroni]* = 0.0032). At 35-40 hpi, SCRs increased in HbAA cultures (median 35.50%) but remained lower in HbSS (median 18.40%, OR[95% CI] = 0.41 [0.27−0.61], *p[Bonferroni]* < 0.0001) and HbSC cultures (median 11.90%, OR[95% CI] = 0.30 [0.14−0.64], *p[Bonferroni]* = 0.0003) (S3B and S3C Figs, S3 Table). While the reduced SCRs observed in HbSS and HbSC RBCs under both basal and inducing conditions may reflect impaired SC associated with these HBB genotypes, it is also possible that these results are explained by parasite developmental arrest in these RBCs. On the other hand, in choline-depleted cultures, SCRs were similar in HbAS or HbAC RBCs compared with HbAA RBCs (Figs S3B and S3C, S1B and S3 Tables), indicating that the high SCR observed after choline depletion does not further increase in mutant HBB RBCs.

The final total parasitemia (Generation 1) followed a similar trend as final gametocytemia between all genotypes, highest in cultures with HbAS RBCs and lowest in HbSS and HbSC RBCs (Fig 3C, S3D Fig). This indicates successful reinvasion of all RBC types, and confirms that the higher SCRs in HbAS reflect enhanced gametocyte production rather than reduced asexual parasite numbers. Furthermore, differences in SCRs could not be explained by variation in initial parasitemia at 0-5 hpi (Generation 0, all cultures were adjusted to a parasitemia below 2.0%), as no significant differences were found between HBB genotypes (beta-binomial GLM, *p [Bonferroni]* = 1; S1B Table). Additionally, no correlation was observed between SCR and initial parasitemia (Spearman ρ = 0.10, *p* = 0.63; S1B Fig). We assessed other possible confounders for the differences in SCR among HBB genotypes. A risk analysis of different variables in the study population (HBB genotype, ABO blood group, gender, village and ethnicity) confirmed that HBB genotypes are a risk factor for enhanced SCRs, in addition to gender (females) (Fisher’s exact test, overall *p* < 0.01 and *p* < 0.05, respectively; Table 1). Gender was not included in further analyses due to its imbalanced spread across HBB genotypes (S4 Table). Since reticulocytes have been proposed to enhance SCRs [63], and individuals with mutant HBB genotypes often exhibit anemia-associated reticulocytosis, we assessed reticulocyte percentages across all HBB genotypes. We observed significantly higher reticulocyte percentages in HbSS compared to HbAA RBCs (beta-binomial GLM, OR [95%CI] = 2.56 [1.20−5.44], *p [Bonferroni]* = 0.008, S1B Table and Fig 3D), which was confirmed in a risk analysis of reticulocytes, amongst other factors, across HBB genotype groups (Kruskal-Wallis test, overall *p* < 0.001; S4 Table). SCRs across HBB genotypes were negatively correlated with reticulocyte percentage (Spearman ρ = −0.50, *p* < 0.01; Table 1). Since SCR could not be readily measured in HbSS and HbSC genotypes due to parasite developmental delay/arrest in these RBCs (see next section), we performed correlation analysis without the HbSS and HbSC genotypes, only between HbAA, HbAC and HbAS. No correlation was observed between reticulocyte percentage and SCR (Spearman ρ = 0.10, *p* = 0.73; S3E Fig), suggesting that higher SCRs are likely due to intrinsic properties of mutant RBCs rather than to higher reticulocyte abundance. Lastly, to control for hemolysis during the assays, we assessed RBC counts (RBCs/µl) across HBB genotypes at various timepoints, as shown in S5 Table. No significant differences were observed at any of the time points, indicating that mutant HBB RBCs did not undergo hemolysis during the course of the experiment.

**Table 1.** Risk analysis of factors associated with elevated SCRs in the *in vitro* SC assays.

| Median (IQR) | SCR at 35-40 hpi (Generation 1) | p-value<sup>§</sup> |
| --- | --- | --- |
| **HBB genotype** |  | <0.01 |
| HbAA | 10.8 (0.49) |  |
| HbAC | 13.30 (4.71) |  |
| HbAS | 16.50 (5.64) |  |
| HbSS | 6.96 (3.17) |  |
| HbSC | 7.99 (1.83) |  |
| **ABO blood group** |  | 0.15 |
| O+ | 7.57 (7.86) |  |
| A+ | 8.80 (2.52) |  |
| B+ | 17.80 (3.48) |  |
| A- | 9.34 (2.87) |  |
| B- | 10.90 (3.77) |  |
| **Gender** |  | <0.05 |
| Female | 11.30 (3.75) |  |
| Male | 7.57 (4.09) |  |
| **Village** |  | 0.30 |
| Nanoro | 11.30 (3.75) |  |
| Nazoanga | 15.40 (0.60) |  |
| Soum | 0 (0.00) |  |
| Séguédin | 0 (0.00) |  |
| **Ethnicity** |  | 0.40 |
| Gourounsi | 17.80 (0.00) |  |
| Mossi | 12.30 (4.14) |  |
|  | **Median (IQR)** | **p-value (rho)<sup>°</sup>** |
| **Reticulocytes (%)** | 1.50 (5.63) | < 0.01 (-0.50) |
| **Age** | 27.00 (30.00) | 0.37 (-0.18) |
| **n (%)** | 28 (100.00) | - |

Significant p-values ( $p < 0.05^*$ , $p < 0.01^{**}$ , $p < 0.001^{***}$ ). §Categorical variables test: Fisher's exact test, Continuous variables test: Kruskal Wallis test, °Continuous variables: Spearman correlation test.
SCRs measured at 35-40 hpi (Generation 1). Data from *P. falciparum* non-infected samples collected in Nanoro, Burkina Faso and the Antwerp University Hospital (UZA, Belgium) (total $n = 34$ ). Statistical tests performed are shown at the bottom of the figure with correspondent significant p-values as: $p < 0.05^*$ , $p < 0.01^{**}$ , $p < 0.001^{***}$ . **Abbreviations:** Hb, hemoglobin; HBB, human hemoglobin beta; hpi, hours post-invasion; IQR, Interquartile range; *ivSCAs*, *in vitro* sexual conversion assays; SCRs, sexual conversion rates.

### Red blood cells with mutant HBB genotypes delay parasite development

*P. falciparum* growing in HbAS RBCs exhibits developmental delay/arrest before DNA replication when growing under low oxygen conditions (≤ 5% O₂, as present in the bone marrow and spleen, and in standard *P. falciparum* cultures as used for the *iv*SCA) [25].

Developmental delay/arrest in a particular HBB genotype would affect the time at which SC can be measured *in vitro.* To characterize in more detail the possible alterations in parasite development when grown in RBCs with different genotypes, we first quantified the percentage of rings, trophozoites and early/late schizonts in a set of cultures with the different HBB genotypes using light microscopy (LM) at three timepoints (35-40 hpi at Generation 0; 25-30 hpi at Generation 1; and 35-40 hpi at Generation 1) (Fig 4A). Parallel analysis of the proportion of late stages (schizonts) in the same samples by flow cytometry revealed consistent patterns between the two approaches (Fig 4B). We therefore used flow cytometry (gating strategy in Fig 4C) to analyze parasite development in all RBC samples. Samples with very low late-stage proportions (at 35-40 hpi) likely reflect slow development (remaining in early stages), rather than rapid development to ring stages in the next generation.

**Fig 4.**
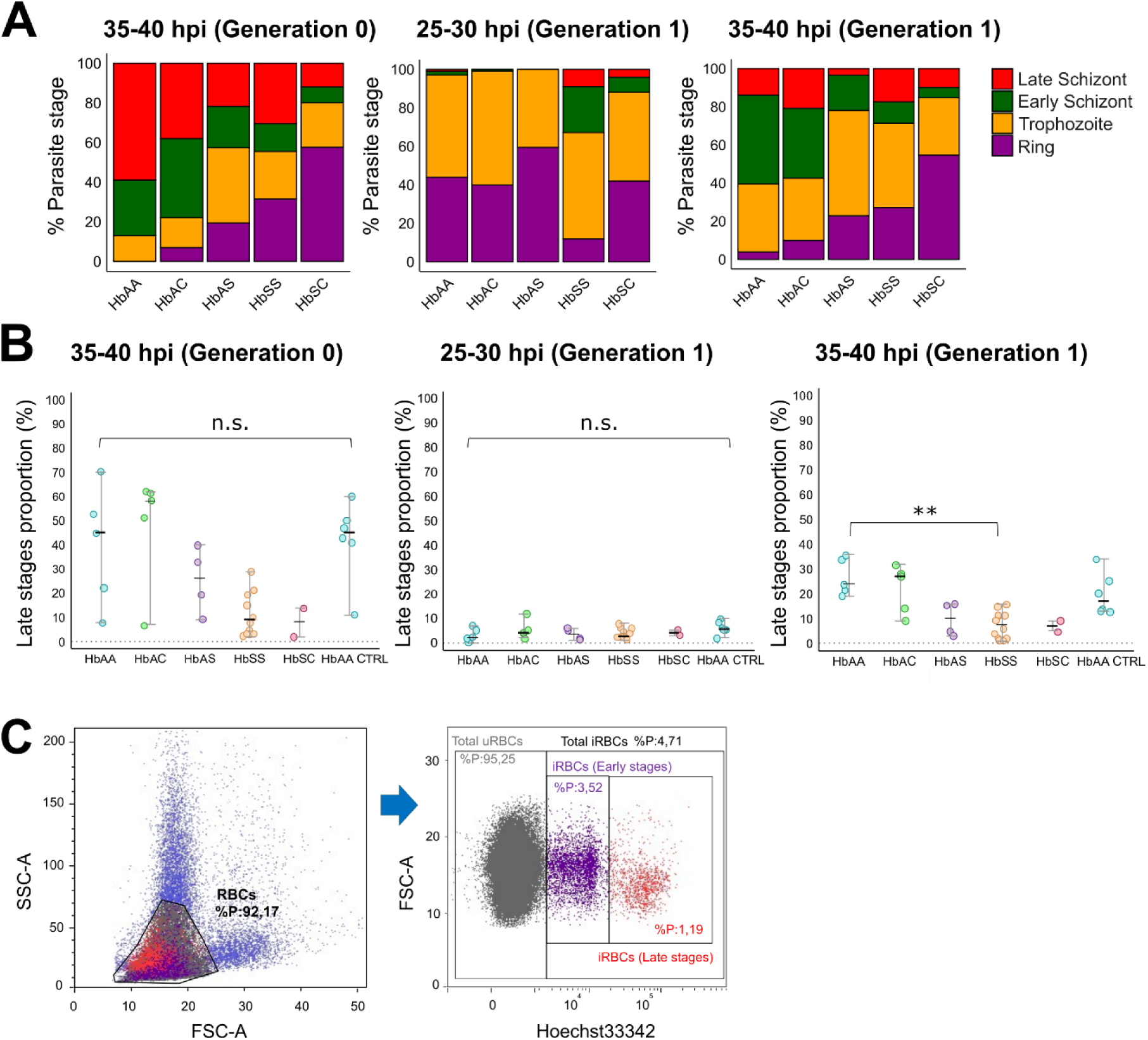
*P. falciparum* parasite development in RBCs with different HBB genotypes. **(A-B)** Timepoints shown are: 35-40 hpi (Generation 0; left graph), 25-30 hpi (Generation 1; middle graph) and 35-40 hpi (Generation 1; right graph). **(A)** Microscopy-based quantification of parasite stages expressed as percentages of a total of 100 counted parasites per sample. Y-axis: percentage of parasite stages (%); X-axis: HBB genotype groups (HbAA *n* = 1, HbAC *n* = 1 and HbAS *n* = 2, and HbSS *n* = 2 and HbSC *n* =2 donors from a malaria-endemic area in Nanoro, Burkina Faso. **(B)** Proportion of late stages (*i.e.,* schizont stages) on the total infected RBCs (iRBCs) by flow cytometry (Y-axis) by HBB genotype groups (X-axis) (HbAA *n* = 5, HbAC *n* = 5 and HbAS *n* = 4, and HbSC *n* = 1 donors from a malaria-endemic area in Nanoro, Burkina Faso; HbSS *n* = 12, HbSC *n* = 1 and *n* = 6 control HbAA donors [HbAA CTRL] from a non-endemic area). **(C)** Representative flow cytometry plots showing the gating strategy for the detection of late parasite stages. Left: total RBCs gated. Right: within the total RBCs gate in left plot, uninfected RBCs (uRBCs) and iRBCs are identified, with iRBCs further gated into early and late parasite stages by Hoechst 33342. **Statistical tests**:(B, right graph) beta-binomial GLM with Bonferroni correction for pairwise comparisons; (B, left and middle graph) Kruskal-Wallis test. Significant (*p* < 0.05*, *p* < 0.01**, *p* < 0.001***) and non-significant *p*-values (ns). **Abbreviations:** CTRL, control sample; FSC-A, Forward Scatter-Area; GLMs, Generalized linear models; Hb, hemoglobin; HBB, human hemoglobin beta; Hpi, hours post-invasion; RBCs, red blood cells; SSC-A, Side Scatter-Area.

Parasite developmental delay was estimated by comparing parasite stage distributions and mean parasite age to wild-type HbAA cultures, using the expected timing of the intraerythrocytic cycle (∼44 h) to approximate the corresponding delay in hours. Delay versus (partial) arrest was defined based on consistent deviations from HbAA cultures across multiple LM- and flow cytometry-derived parameters, including late-stage proportions at successive time points and their change between generations (Fig 3A), as well as parasite multiplication rates, allowing classification of individual cultures as expected, delayed, or (partially) arrested/extremely delayed development (S6 Table). Detailed analysis is further described in *S1 Appendix.* We found that parasites in HbAA RBCs progressed normally from Generation 0 to 1 with no apparent delay; however, there was unexpected high variability between individual donors (Fig 4B). In HbAC RBCs, parasite development was slightly delayed (∼3 hours delay vs. HbAA at Generation 0, 35-40 hpi) but without arrest. In contrast, HbAS RBCs showed a more pronounced delay (∼10 hours), with partial developmental arrest observed in one sample (Figs 4A and 4B, S6 Table). HbSS and HbSC RBCs presented a similar delay to HbAS (∼9 and ∼16 hours delay, respectively) with a persistent population of schizonts, trophozoites, and late rings across all timepoints (Fig 4A), consistent with the high frequency of partially arrested/delayed cultures (67% in HbSS and 50% in HbSC) (S6 Table). The (partial) arrest/extreme delay in these cultures potentially makes them unsuitable for SCR measurement and may explain the apparent reduction in SCRs associated with growth in RBCs with these genotypes. These results confirm the convenience of measuring SCRs at 35-40 hpi in this experimental set-up.

### IgG and IgM reactivity in plasma of individuals with different HBB genotypes

We analyzed IgG and IgM reactivity against *P. falciparum* in individuals with different HBB genotypes using flow cytometry and selected plasma samples from humans living in the malaria-endemic area of Nanoro, Burkina Faso, collected as part of the *InHost* longitudinal cohort study [74] (see *Materials and methods*). IgG and IgM antibodies were measured against antigens on the surface of intact RBCs infected with the *P. falciparum* transgenic reporter line NF54-*gexp02-Tom* [65, 81] at the early trophozoite / gametocyte stage (20-25 hpi; Generation 1; Fig 3A) using a flow cytometry protocol adapted from Dantzler KW., et al., (2019) [82]. Plasma samples from individuals living in malaria-endemic area were defined as “endemic plasma”, whereas samples from individuals living in non-endemic area were defined as “non-endemic plasma”. Detailed information on assay optimization and validation is provided in the *S1 Appendix.* Flow cytometry gates were set to quantify total iRBCs (trophozoites and stage I gametocytes), asexual iRBCs (trophozoites), and sexual iRBCs (stage I gametocytes) based on Hoechst33342 and tdTomato (tdTom) fluorescence. IgG- and IgM-positive cells were identified using AF488 and PerCP-Cy5.5, respectively, resulting in six populations: total iRBCs (Hoechst+/tdTom+ or -), asexual iRBCs (Hoechst+/tdTom-), and sexual iRBCs (Hoechst+/tdTom+), each subdivided by IgG- or IgM-positivity (S4B Fig). To ensure robust ΔMean Fluorescence Intensity (ΔMFI) measurements, a minimum number of 200 IgG- or IgM-positive iRBCs (events) per gated population was required. This cut-off refers to the number of antibody-positive cells contributing to the ΔMFI calculation (Figs S5A and S5C).

In each assay, positive controls including three acute *P. falciparum*-infected non-endemic plasma samples from a traveler at days 0 (D0), 21 (D21) and 23 (D23), and a negative control (“naïve non-endemic plasma”) were analyzed. Endemic plasma samples were considered positive for IgG or IgM when both the event count and IgG- and IgM-positive total-iRBC proportions exceeded those observed in the “naïve non-endemic plasma”. The “naïve non-endemic plasma” sample consistently did not reach the 200 event counts threshold, confirming its suitability as a negative control. The negative control values served as a baseline for the IgG- and IgM-positive total iRBC proportions (% cells) and were lower than those of the endemic plasma samples of all HBB genotypes (Fig 5). Importantly, the events count of IgM-positive total and asexual iRBCs in the positive controls at D21 and D23 exceeded both the event counts observed in the “naïve non-endemic plasma” and the threshold of 200 events, confirming that our flow cytometry assay can detect IgM-positive total and asexual iRBCs (S5C Fig). The time course of the “acute *P. falciparum*-infected non-endemic plasma at D0, D21 and D23” generally followed the expected IgG (D0 < D21 < D23) and IgM (D0 < D21 > D23) dynamics in IgG- and IgM-positive total iRBCs after a *P. falciparum* infection [83, 84].

**Fig 5.**
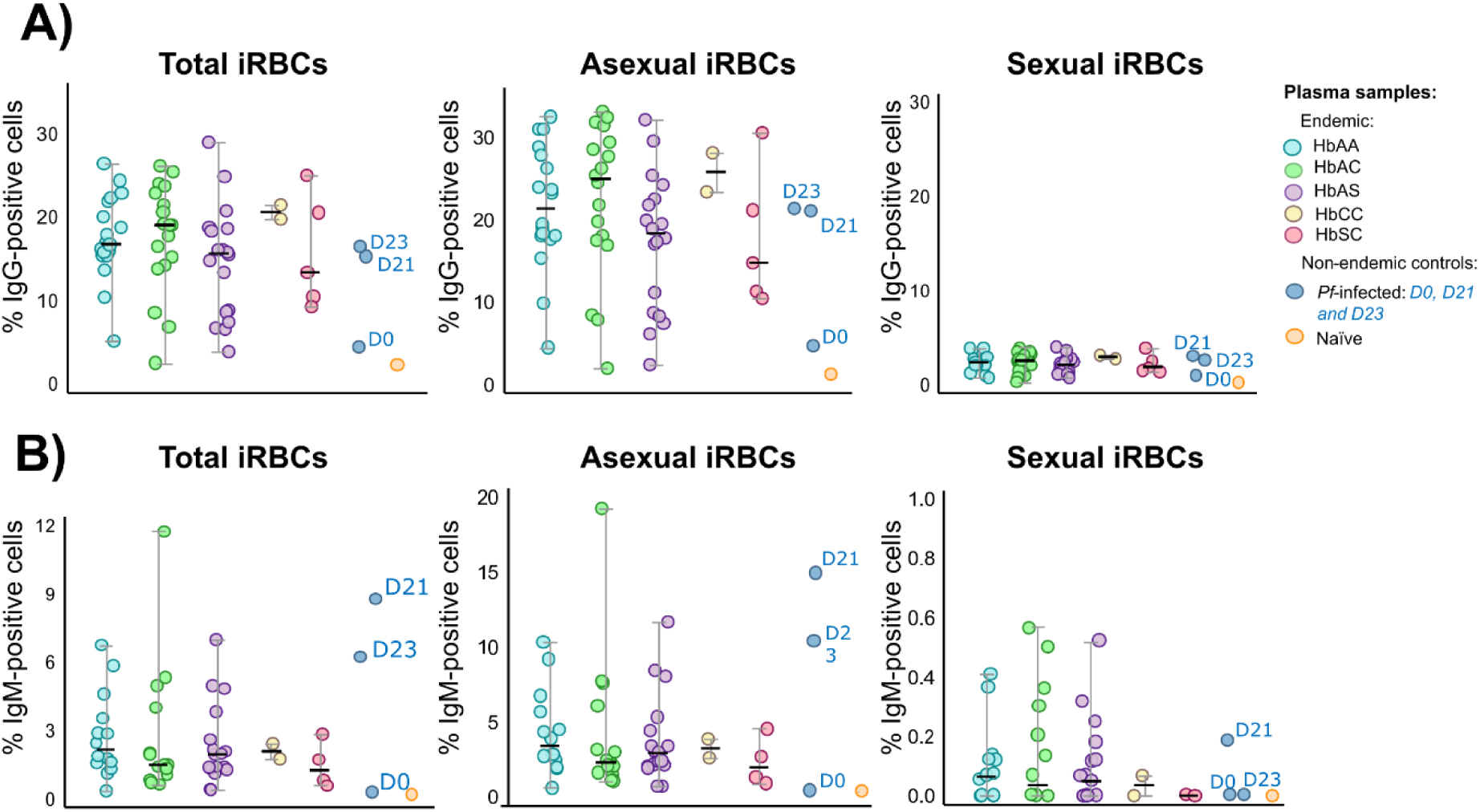
IgG- and IgM-mediated responses to *P. falciparum* trophozoites and stage I gametocytes across HBB genotypes. Data from *P. falciparum* infected (*n* = 44) and non-infected (*n* = 17) plasma samples collected at the Clinical Research Unit (CRUN) in Nanoro, Burkina Faso (total *n* = 61). **(A)** Percentage of IgG-positive iRBCs: total (left), asexual (middle) and sexual (right) iRBCs. **(B)** Percentage IgM-positive iRBCs: total (left), asexual (middle) and sexual (right) iRBCs. Positive controls (“*Pf*-infected non-endemic” *i.e.,* three acute *Pf-*infected non-endemic plasma samples from a traveler at D0, D21, and D23) and a negative control *i.e.,* “Naïve non-endemic”: non-endemic plasma from a volunteer with HbAA genotype) are shown as individual dots per sample. The Y-axis range differs between graphs for visualization. No significant differences were found (beta-binomial GLM with Bonferroni adjustment for pairwise comparisons, *p* > 0.05). **Abbreviations:** asexual iRBCs, trophozoites; D, day(s); GLM, Generalized linear models; Hb, hemoglobin; HBB, human hemoglobin beta; IgG and IgM, Immunoglobulin G and M; iRBCs, infected-red blood cells; *Pf, P. falciparum*; sexual iRBCs, stage I gametocytes; total iRBCs, trophozoites and stage I gametocytes.

MFI could only be evaluated for IgG-positive total and asexual iRBCs in the statistical analyses (Figs S5B and S5D), because all endemic plasma samples (*n* = 61) exhibited event counts below 200 for the IgM and IgG-positive sexual iRBCs, thirteen samples (21%) had no detected (*i.e.,* zero) events for IgM-positive sexual iRBCs and the median event count of IgM-positive total and asexual iRBCs was below the 200 event threshold for all HBB genotype groups. The percentage of IgG- or IgM-positive iRBCs reflects the proportion of iRBCs recognized by antibodies, whereas the MFI reflects the relative level of antibody binding per positive cell. When IgM-positive sexual iRBCs were rare or absent, the proportion of positive cells could still be quantified, but MFI could not be reliably estimated due to insufficient antibody-positive events.

In endemic plasma samples, we detect low or absent IgG- and IgM-positive sexual iRBC proportions, and low or absent IgM-positive total and asexual iRBC proportions, as compared to high IgG-positive total and asexual iRBC proportions (Figs 5 and S5). We found no significant differences in the IgG- and IgM-positive total iRBC proportions (% of cells/ΔMFI data are only considered robust for IgG-positive total and asexual iRBCs, reaching the minimum threshold of 200 events; Fig 5) (beta-binomial GLM, *p* > 0.05; S1C Table). When analyzing the response against sexual or asexual iRBCs separately, there was no significant difference across endemic plasma from individuals carrying different HBB genotypes in IgG- or IgM-positive asexual or sexual iRBC proportions (Fig 5, S7 Table) (beta-binomial GLM, *p* > 0.05; S1C Table).

The evaluation of variables in the study population (*n* = 61) such as: *P. falciparum* infection status at the time of sample collection, parasitemia (parasites per µl), presence of gametocytes in blood, age, gender and village; across HBB genotype groups (HbAA [*n* = 18], HbAC [*n* = 18], HbAS [*n* = 18], HbCC [*n* = 2], and HbSC [*n* = 5]) revealed no significant differences between categories (Fisher’s exact test and Kruskal-Wallis test; *p* > 0.05; S8 Table). In our study population, there is a homogeneous distribution of individuals across the categorical variables, with no observed sampling bias that could affect further analysis (S8 Table).

Further analysis of the IgG- and IgM-positive total iRBC proportions by “*P. falciparum* infection status”, showed higher antibody recognition in infected individuals (*n* = 44) compared to non-infected individuals (*n* = 17) (S6 Fig). IgG-positive total iRBC proportions were markedly higher in infected individuals (median[IQR]: 17.90[6.30]) than in non-infected individuals (9.10[9.66]) (beta-binomial GLM, OR[95%CI] = 1.81[1.35−2.43], *p* = 0.001; S1C Table). In contrast, IgM-positive total iRBC proportions showed only a modest increase in infected individuals (2.03[3.34]) compared to non-infected individuals (1.56[1.27]) (beta-binomial GLM, OR[95%CI] = 1.51[1.01−2.23], *p* = 0.043; S1C Table). These results confirm the expected IgM dynamics, where IgM-positive iRBC proportions are highest during acute or recent infections (*P. falciparum*-infected individuals) compared to past infections (non-infected individuals) [83, 84].

Stratification analysis of the IgG- and IgM-positive total iRBC proportions by age-group (*i.e.,* younger age: “1 to 4 years” compared to older age: “5 to 19 years”) in *P. falciparum*-infected compared to non-infected individuals, showed that both *P. falciparum*-infected and non-infected individuals with younger age had lower IgG-positive total iRBC proportions (*Pf*-infected *n* = 22 and median[IQR]: 17.90[7.23]; non-infected *n* = 15 and median[IQR]: 9.30[10.80]) than those with an older age (*P. falciparum*-infected *n* = 22 and median [IQR]:18.60[5.64]; non-infected *n* = 2 and median[IQR]: 12.60[3.69]) (Welch’s ANOVA, *p* = 0.082 [IgG]; Kruskal-Wallis test, *p* = 0.38 [IgM]). *P. falciparum*-infected individuals with younger age had slightly higher IgM-positive total iRBC proportions (*n* = 22, median [IQR]: 2.19[2.33]) than those with an older age (*n* = 22, median [IQR]: 1.87[3.04]) (Kruskal-Wallis test, *p* = 0.94). In contrast, the IgM-positive total iRBC proportions in non-infected individuals with younger age were slightly lower (*n* = 15, median[IQR]: 1.56[1.12]) than those with an older age (*n* = 2, median [IQR]: 2.18[0.00]) (Kruskal-Wallis test, *p* = 0.16).

Overall, these findings demonstrate that the flow cytometry assay accurately detects IgM-positive asexual iRBC proportions, as validated by positive controls, whereas such responses are rare or absent in individuals from endemic areas (Fig 5). The increase of IgG with age and decrease of IgM with age explains the low IgM-positive total iRBC proportions in the endemic plasma: exposure leads to a build-up of immunity that replaces low-affinity IgM by an affinity-matured IgG response [84–88].

## DISCUSSION

This study demonstrates that *P. falciparum* SC is significantly influenced by host hemoglobin genotype. SC was higher in HbAS RBCs than in wild-type HbAA RBCs both in natural infections *in vivo* and *in vitro* (using the *P. falciparum* transgenic reporter line NF54*-gexp02-Tom*), whereas a slight increase was observed for HbAC RBCs *in vivo* that could not be confirmed *in vitro*. This genotype-dependent modulation of SC was accompanied by a developmental delay in parasites within HbS- and HbC-infected RBCs (iRBCs). However, we found no differences in IgG or IgM responses against trophozoite- and early gametocyte-iRBC surface antigens across HBB genotypes. Together, these findings indicate that host RBC intrinsic properties, rather than humoral immunity, underly altered parasite reproductive strategies in HBB mutant backgrounds.

Amoah et al., (2020) [89] analyzed the effect of HBB genotype on SC of field isolates from Ghanaian children aged 6-15 years with uncomplicated malaria, and reported a non-significant trend of decreasing SCRs from HbAA to HbAC and HbAS. The discrepancy with our results is most likely methodological, because quantifying SC in natural infections is challenging as parasites in freshly collected blood include morphologically indistinguishable sexual and asexual rings. The authors used prolonged culture (≥4 days) to distinguish sexual stages from asexual forms, similarly to Usui et al., (2019) [49]. The limitation of this approach is that SCRs are measured as the ratio of stage II gametocytes on day 4 (D4) by the total parasitemia on day 0 (D0). This can underestimate SCR because parasites contributing to the denominator (D0) are counted before culture adaptation affects viability, whereas the numerator (D4) includes only gametocytes that survive and mature after several days in *ex vivo* conditions, during which parasite loss can occur.

We addressed these limitations by adapting an *ev*SCA to quantify SC [45] for experiments with mutant RBCs. We measured whether infection of individuals with RBCs with different HBB genotypes leads to alterations of sexual conversion rates. We quantified the proportion of parasites in patient samples that differentiated as asexual or sexual forms after *ex vivo* maturation. Quantification is done by immunofluorescence, with stage I gametocytes detected by Pfs16 IFA and overall parasites by DAPI [80]. Key aspects for the success of our new *ev*SCA were the inclusion of the PKG inhibitor ML10 that prevented reinvasion, allowing numerator and denominator measurements for the SCR at the same time point, while enrichment using LD columns improved sensitivity. This approach reliably quantified SC in natural human infections with densities as low as 0.05% (∼2,500 parasites/µl) and detected a minimum of 38 stage I gametocytes per sample, with no gametocyte-negative cultures.

We confirmed the *in vivo* findings in the *ev*SCAs using an *in vitro* SC assay (*iv*SCA) with a gametocyte reporter line [81], which allows early detection of sexual rings. In this assay, parasites growing in HbAS RBCs showed higher SCRs than those in HbAA RBCs, whereas parasites in HbSS and HbSC RBCs were associated with lower SCRs *in vitro*. For HbAC RBCs, a slight increase observed although not statistically significant. The fold-change SCR observed for HbAS relative to HbAA RBCs in the *iv*SCA was consistent with that reported by Flage, B., et al., (2023) [32] (1.5 vs. 1.5 in HbAS), although the results are not directly comparable due to clear methodological differences. In our assay, SCR was defined as the proportion of *gexp02*-expressing parasites among total parasites in the next cycle (generation 1), whereas in Flage, B., et al., (2023) [32], sexual parasites were quantified at generation 1 and total parasitemia at generation 0, precluding direct comparison. For HbSS and HbSC RBCs, strong conclusions regarding SCR cannot be drawn, as parasite development appeared to be impaired after reinvasion and no corresponding data from natural infections were available. This contrasts with findings in Flage, B., et al., (2023) [32], who reported markedly higher SCRs in HbSS RBCs (35% vs. 6.96% in our study) and no apparent developmental delay at day 2 of culture, although this was not quantitatively assessed. As noted above, differences in the timing and approach used to quantify SCR make direct comparison between studies difficult.

Parasite developmental progression differed across HBB genotypes and likely influenced SCR estimation. We observed a partial developmental delay in parasites infecting HbAS and HbAC RBCs, and a more pronounced arrest or extreme delay in HbSS and HbSC RBCs under low-oxygen conditions (5% O₂), as assessed by both light microscopy and flow cytometry, consistent with Archer, N.M., et al., (2018) [25]. Later time-point measurements at 35-40 hpi resulted in higher SCRs in endemic HbAA, HbAC, and HbAS RBCs compared with 25-30 hpi, suggesting delayed sexual marker expression and underestimation of SCR at earlier time points due to developmental delay. In contrast, parasites grown in control non-endemic HbAA RBCs showed stable SCRs across both time points. The differences observed between endemic and non-endemic HbAA RBCs may reflect unmeasured host genetic factors, such as blood group variants or other RBC polymorphisms.

Under choline-depleted conditions, SCRs in HbSS and HbSC increased modestly (from <10% to 10-20%) but remained below levels in HbAA, HbAC, and HbAS RBCs (>30-40%). This likely reflects severe developmental delay causing under-detection of sexually committed rings at the measured time points rather than inhibited sexual commitment *per se*. Overall, our data indicate that accurately assessing SC in HbSS and HbSC RBCs is challenging with this experimental approach due to impaired parasite development. We further evaluated host-related factors previously associated with SC, including reticulocyte abundance, parasitemia, and ABO blood group [63–65, 90–94], but none of these potential confounders influenced SCRs in either the *ev*SCA or *iv*SCA. Together, these results indicate that the observed increase in SC is specifically associated with HBB genotype rather than these other host-related variables. Interpretation of risk/GL(M)M analyses was limited by the small sample size and unbalanced group structure in terms of potential confounding factors, which constrained model complexity and precluded fully adjusted multivariable analyses. Although residual confounding or interactions cannot be excluded, the direction and magnitude of the observed HBB genotype effect were largely consistent across models, supporting its biological relevance.

The increase in SC in RBCs with HBB mutant genotypes *in vitro*, in the absence of plasma and white blood cells, points to RBC-intrinsic effects rather than host immune responses or soluble plasma factors as the most plausible cause for the altered SCR observed in HBB mutant RBCs. HbS and HbC RBCs exhibit elevated levels of free heme [95], linked to increased oxidative stress that leads to polymerization of HbS under low oxygen conditions [96], and delays parasite development [25]. In Flage, B., et al., (2023) [32], authors showed that heme directly induces SC. The oxidative stress inhibits actin polymerization in infected HbS and HbC RBCs [34], which alters protein trafficking and PfEMP1 display [28, 31]. This reduces cytoadherence [35], and membrane flexibility [97]. Oxidative stress also modifies RBC micro RNAs that can hybridize with parasite mRNAs [27], directly affecting parasite development and metabolism, and increasing gametogenesis [32]. Taken together, these findings suggest that the effect of HbAC and HbAS RBCs on SC *in vitro* is likely driven by oxidative stress caused by free heme within these RBCs, mediated by their specific miRNA profiles. The stronger effect *in vivo*, particularly in HbAS (7.07-fold vs ∼2.05-fold *in vitro*), probably reflects additional *in vivo* factors such as parasite genetic diversity [49, 80], host factors not captured *in vitro* [80, 98], although we cannot exclude that part of the difference may be attributable to variations in experimental setup.

To further investigate potential *in vivo* influences, we assessed the humoral immune response in the different HBB genotypes. Altered antigen presentation, cytoadherence, and increased splenic clearance [35] could plausibly lead to an modified immune response in HbAC and HbAS. However, epidemiological studies examining immunity in individuals with HbAS and HbAC genotypes have yielded conflicting results. Older children with HbAS have been shown to be more protected against the establishment of parasitemia, suggesting an acquired mechanism of protection [29]. In contrast, another study found no differences in antibody responses between individuals with HbAA, HbAS, and HbAC genotypes against their panel of 491 immunoreactive *P. falciparum* antigens [99]. More recent studies have shown that serological responses tend to be differential in individuals with HbAC and HbAS when focusing on specific surface antigens and populations [70–73].

Given that differences in serological responses appear specific to certain surface antigens, we chose to measure IgG and IgM reactivity against intact iRBCs. This approach enables the assessment of immune responses to native surface antigens, minimizing potential artifacts arising from non-native protein folding, glycosylation, or conformation. We focused on trophozoite-iRBCs (25-30 hpi), as surface antigen presentation (*i.e.,* PfEMP1 expression) begins around 16 hpi and plateaus at approximately 24 hpi [100]. Moreover, commitment to SC occurs between 17-37 hpi [45, 65], suggesting that any antibody-mediated induction of sexual commitment would need to occur at this stage.

We compared IgG and IgM responses to trophozoite- and gametocyte-iRBCs using plasma samples from Burkinabé individuals carrying HbAA, HbAC, HbAS, HbCC, and HbSC genotypes, assessed by flow cytometry. Our analyses revealed no significant differences in IgG or IgM reactivity against trophozoite- and stage I gametocyte-iRBCs across genotypes. IgM responses, as well as responses against gametocyte-iRBCs, were generally low or absent. Therefore, we could not confirm genotype-dependent differences in humoral responses to these parasite stages and consider it unlikely that antibodies directly drive the differential SC observed in these individuals. However, our analysis was limited to the antigens expressed by the parasite reporter line used in the assay. Responses against the immunodominant variant surface antigen PfEMP1 are notoriously difficult to decipher in *in vitro* systems as it is encoded by approximately 60 different *var* genes which are expressed mutual exclusively [101] and expression is significantly reduced *in vitro* compared to *in vivo* [102]. Moreover, we cannot exclude the possibility that genotype-specific antibody responses to later parasite stages, particularly during gametocytogenesis, as suggested by recent studies on gametocyte antigenicity [82, 103], could influence transmission potential, either by promoting clearance of developing gametocytes in the human host or by inducing gametocyte sterility within the mosquito vector [104].

Overall, this study demonstrates elevated SC of *P. falciparum* parasites in HbAS and HbAC RBCs both *in vitro* and in *ev*SCAs − measuring SC in natural human infections *in vivo*. For HbAC RBCs, however, the results were less clear: differences in SC were smaller in both natural infections and *in vitro*, and the *in vitro* differences were not statistically significant, indicating the need of additional studies. These findings further support the view that *P. falciparum* can adjust its reproductive investment in response to stress-related cues. In mutant HBB RBCs, elevated levels of free heme can induce developmental delay and thereby compromise survival within the human host; increased investment in SC may thus represent a parasite survival strategy [25, 32, 105].

Previous studies have shown that microenvironmental factors such as LysoPC depletion, nutrient limitation and exposure to antimalarial drugs can modulate malaria transmission by changing the parasite’s SCR. How factors such as free heme translate into a signal received by epigenetic regulators controlling the sexual commitment genes *ap2-g* or *gdv1* remains an open question [32, 45]. At the population level, HBB mutant genotypes have been linked to increased gametocyte carriage and higher mosquito infectivity. In Burkina Faso, a two- to four-fold higher infection rates in HbAS, HbAC, and HbCC carriers was reported [36]. Similar trends were observed in Mali and Cameroon, where HbAC and HbAS individuals carried more gametocytes and were more infectious to mosquitoes than HbAA [37, 40]. These population-level associations are consistent with our findings, and suggest that increased SC in mutant HBB RBCs may contribute to their enhanced transmission potential.

## MATERIALS AND METHODS

Details about reagents and tools are available in S9 Table.

### Study design

Sample collection for the present study was embedded in a longitudinal cohort study assessing the effect of host factors on transmission potential (*InHost* study, ethical advice reference 1261/18 and 19/06/064), conducted in the Nanoro health district, a rural malaria-hyperendemic region, in the Central-West of Burkina Faso (Fig 6)[74]. Briefly, 871 study participants of all ages from four villages (Nanoro, Nazoanga, Séguédin and Soum) of the district were followed-up for two years (March 2019 - February 2021) through six cross-sectional surveys conducted during both high and low transmission season to assess asexual and sexual parasite carriage and risk factors. Active case detection was complemented by passive case detection at peripheral health centers.

**Fig 6.**
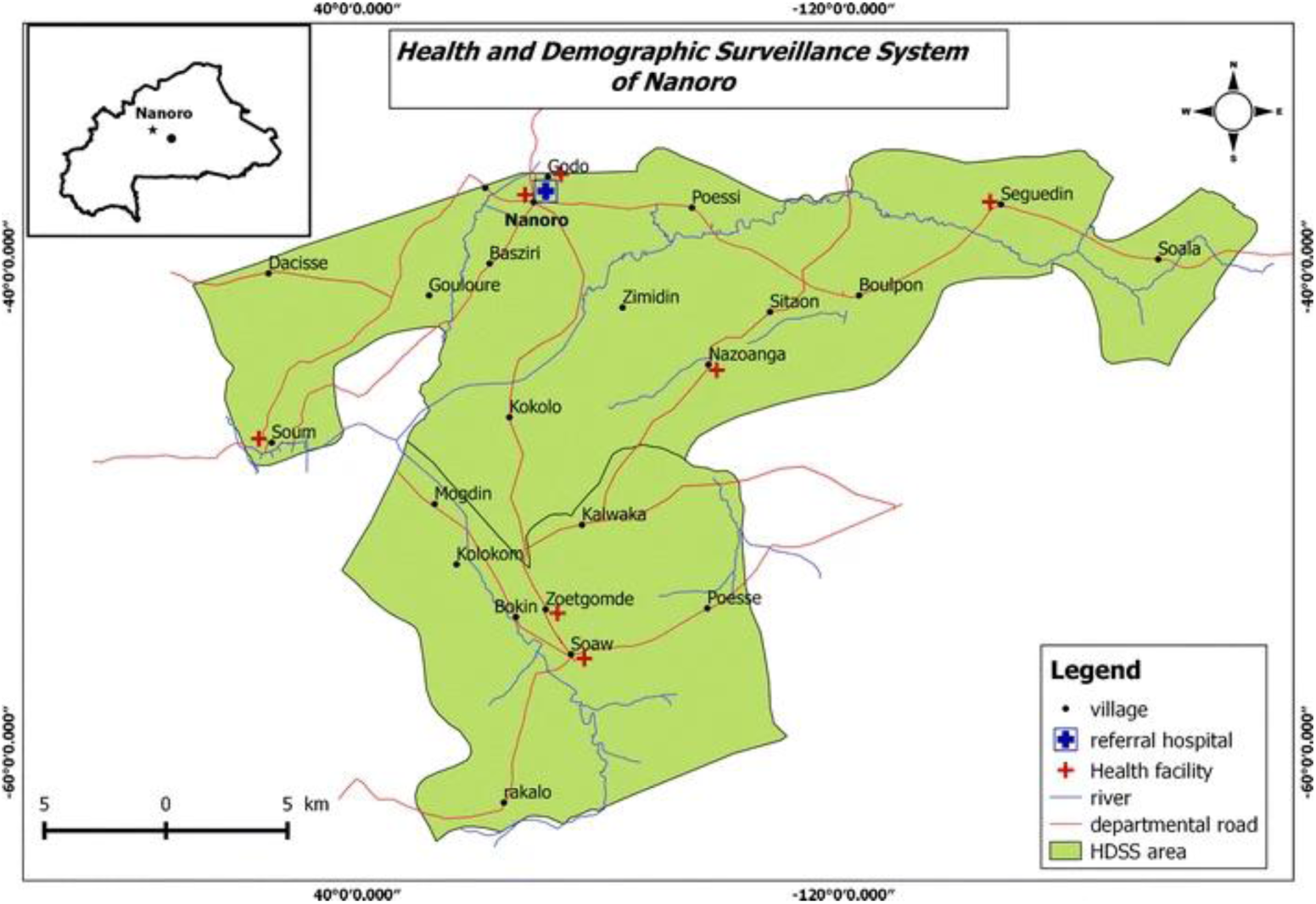
Map of the study area. The inset map highlights the location of the Nanoro health district of Boulkiemdé Province in Central Burkina Faso, with the black dot indicating the position of the capital, Ouagadougou.

For this study, from all participants of the *InHost* study (*n* = 871), 864 individuals were genotyped for their human HBB genotype (see section on “Hemoglobin beta sequencing”). Participants were then selected for this sub-cohort based on their genotype. A venous blood sample was collected from all selected participants, and screened for *P. falciparum* infection. Samples from *P. falciparum-*infected individuals were used to perform *ex vivo* SC assays (*ev*SCAs) and samples from non-infected individuals were preserved to perform *in vitro* SC assays (*iv*SCAs). We included all individuals carrying the HBB genotypes HbAC (*n* = 72), HbAS (*n* = 50), HbCC (*n* = 2), HbSC (*n* = 7), and a random subset of individuals with the HbAA genotype (*n* = 94) (S7 Fig). In total, 225 individuals were listed (age groups “5 - 19 years” [*n* = 166] and “more than 20 years” [*n* = 59]). Listed individuals were visited by a nurse at home and were invited to the Clinical Research Unit of Nanoro (CRUN, Burkina Faso) for screening and enrolment into the study. Population screening for *P. falciparum* infections and enrolment took place over a two-month period (August-September 2022).

At CRUN, the eligibility was assessed based on inclusion and exclusion criteria for the *ev*SCAs (*P. falciparum*-infected individuals) and *iv*SCAs (non-infected individuals). Inclusion criteria consisted of: being enrolled in the main *InHost* study, a confirmed *P. falciparum* malaria infection (for the *ev*SCAs) or a confirmed negative *P. falciparum* malaria infection (for the *iv*SCAs), willing to participate in this study and undergo additional sampling, and signed informed consent. Exclusion criteria consisted of: mixed infection with other *Plasmodium* species (only for *P. falciparum*-infected individuals in the *ev*SCAs), antimalarial treatment within the last two weeks, signs of severe malaria (*e.g.* prostration), severe chronic illness besides hemoglobinopathies (*e.g.,* cancer, heart disease, kidney or liver disorders, HIV/AIDS, etc.), and planning to leave the study region during the study period. Of the 225 listed individuals, 162 were screened at CRUN, of whom 161 participants were selected as eligible and were included within this sub-cohort (S7 Fig). A 15 ml venous blood sample was collected from each participant. After sample collection, blood samples were screened for *P. falciparum* infections (see section on “*P. falciparum* diagnosis”). Samples with confirmed *P. falciparum* infection and parasitemia by light microscopy (LM) ≥ 0.05% (*i.e.* ≥ 2500 parasites/µl) were processed for *ev*SCAs at CRUN (Table 2). Blood samples negative for *P. falciparum* infection were processed and preserved for *iv*SCAs conducted at the Institute of Tropical Medicine (ITM), Antwerp (Table 2, see section on “*iv*SCAs”). *P. falciparum*-infected individuals, as detected by LM and confirmed with *P. falciparum var* gene acidic terminal sequence (*var*ATS) quantitative PCR (qPCR), were treated with Artemether-Lumefantrine (AL) following national guidelines and were asked to return three days later to confirm infection clearance.

**Table 2.** Number of samples (*n*) per *Plasmodium falciparum* infection status and donor HBB genotype at collection. Information given for the *ex vivo* SC assays *(ev*SCAs), *in vitro* SC assays (*iv*SCAs) and Immunoglobulin (Ig) detection assays.

|  | HbAA | HbAC | HbAS | HbSC | HbCC | HbSS | Total |
| --- | --- | --- | --- | --- | --- | --- | --- |
| **EvSCAs** |  |  |  |  |  |  |  |
| *Pf*+ (CRUN, total *n*) | 14 | 12 | 6 | - | - | - | 32 |
| **ivSCAs** |  |  |  |  |  |  |  |
| *Pf*- (total *n*) | 5 | 5 | 4 | 2 | - | 12 | 28 |
| CRUN (*n*) | 5 | 5 | 4 | 1 | - | - |  |
| UZA (*n*) | - | - | - | 1 | - | 12 |  |
| **Ig detection** |  |  |  |  |  |  |  |
| *Pf*+ (CRUN) | 13 | 13 | 13 | 3 | 1 | - | 43 |
| *Pf*- (CRUN) | 5 | 5 | 5 | 2 | 1 | - | 18 |
| Total *n* (*Pf*+ and *Pf*-) | 18 | 18 | 18 | 5 | 2 | - | 61 |

Note: *P. falciparum*-infected (*Pf*+) samples used in the *ev*SCAs must have a parasitemia $\geq 0.05\%$ (*i.e.*, $\geq 2500$ parasites/ μl), as the established limit of accurate assessment of SC with these assays. **Abbreviations:** CRUN, Clinical Research Unit in Nanoro; Hb, hemoglobin; ITM, Institute of Tropical Medicine in Antwerp; *Pf*, *P. falciparum*; UZA, Antwerp University Hospital.

Sample size was calculated based on a power analysis assuming an expected effect size of 50%, a 5% significance level, and 90% power. This sample size was considered sufficient to detect differences in SC by HBB genotype, based on effect sizes reported in previous studies [63]. To account for potential variability in HBB genotype groups, a minimum of 8 participants per HBB genotype group was required. To ensure sufficient sampling and successful assays per HBB genotype group, the sample size was upscaled to a maximum of 15 participants per HBB genotype group.

To increase the number of HbSC and HbSS blood samples for *iv*SCA, additional blood samples were collected from patients with hemoglobinopathies (HbSS and HbSC) visited at the Antwerp University Hospital (UZA), Belgium. A total of 13 adults (aged 19-37 years), routinely followed at UZA’s Hematology Unit, were invited to donate a blood sample with exclusion criteria of fever. The inclusion period was between October 2023 and April 2024.

### Plasmodium falciparum diagnosis

*P. falciparum* malaria diagnosis was done by RDT using the Bioline™ malaria antigen P.f^®^ test (05FK50, Abbott) (recommended by the National malaria Control Programme (NMCP) in Burkina Faso), following the manufacturer instructions. In addition, malaria parasites were detected and quantified by LM following standard protocols at CRUN [106]. In brief, venous blood smears were stained with Giemsa and examined until 200 white blood cells were counted. Parasitemia was calculated based on the number of asexual parasites per white blood cells, expressed as asexual parasites per microliter of blood, assuming a white blood cell count of 8,000 per microliter. The presence of gametocytes and other *Plasmodium* species (*P. ovale and P. malariae*) was assessed in all positive blood smears [107]. Blood samples that tested positive for *P. falciparum* by both RDT and LM were classified as *P. falciparum*-infected. Samples that were negative by LM (regardless of the RDT result), were identified as non-infected (Table 2, S7 Fig). Negative infection with *P. falciparum* was confirmed by *var*ATS qPCR, using five μl of DNA (extracted with QIAamp DNA Mini Kit, Qiagen Germany), as previously described [108, 109] (S7 Fig). The limit of detection of the *var*ATS qPCR was 0.1 parasite/μl (in our hands), with samples showing Ct > 39.7 considered negative. DNA from the *P. falciparum* 3D7 strain served as a positive control, while negative controls included DNA from non-infected human blood and a no-template control (NTC).

### Sample collection and processing

At CRUN, a 15 ml venous blood sample was collected in an Acid Citrate Dextrose (ACD) tube and the residual blood in the needle was used for *P. falciparum* diagnosis by RDT and LM (see section on “*P. falciparum* diagnosis”). The sample was then immediately transported to the laboratory for processing. Samples were centrifuged (1,648 g for 5 minutes) and washed two times with RPMI to isolate the RBC pellet. RBC pellets from *P. falciparum*-infected blood samples were immediately put in culture for the *ev*SCAs (see section on “*Ex vivo* SC assays (*ev*SCAs)”. RBC pellets from non-infected samples were cryopreserved (1:3 dilution in cryopreservation solution containing 3% sorbitol and 0.65% NaCl) and stored directly at −80°C until shipment to ITM, where preserved in liquid nitrogen until use in *iv*SCAs. Two hundred microliters of whole blood were used for DNA extraction and confirmation of *P. falciparum* infection by *var*ATS qPCR (see section on “*P. falciparum* diagnosis). At UZA, 15-20 ml of non-infected venous blood samples were collected in EDTA tubes and immediately transported at 4°C to ITM for processing, done as described at CRUN. The resulting RBC pellets were cryopreserved (1:3 dilution in cryopreservation solution containing 3% sorbitol and 0.65% NaCl) by direct snap-freezing in liquid nitrogen for subsequent *iv*SCAs. Additionally, ABO and Rhesus D blood groups were determined with agglutination tests (BIO-RAD, REF: 801320, 801345, 802032).

### Hemoglobin beta sequencing (HBBseq)

In samples collected at CRUN, PCR amplification of the HBB gene (NCBI_Gene:3043) was performed to identify single nucleotide polymorphisms (SNPs) using PCR conditions adapted from Wittwer CT,. et al., (2020)[110] and MMV & WHO [111]. Primer pairs were designed to amplify a 599 bp product (S9 Table). PCR reactions were performed in a total volume of 25 µl, containing 2.5 µl of DNA template, 10× Buffer B, 5 mM of each dNTP, 0.3 µM of each primer, 25 mM of MgCl₂, and FIREPol® DNA Polymerase (Solis Biodyne, Ref: 01-01-00500). Thermal cycling conditions were as follows: an initial denaturation at 94°C for 6 minutes, followed by 40 cycles of 60.6°C for 30 seconds, and 72°C for 1 minute, with a final extension at 72°C for 10 seconds. After Sanger sequencing, chromatograms were analyzed using BioEdit 7.2. (Informer Technologies) to identify the wild-type and the mutant HBB codons (Table 3).

**Table 3.** HBB codons and genotypes identified by Sanger sequencing analysis of the HBB gene (HBB-seq).

| HBB allele | Mutation | HBB codon |
| --- | --- | --- |
| HbA (wild-type) | - | GAG |
| HbS (mutant) | *c.20A>T*<br>= E (Glu) > V (Val) | GTG |
| HbC (mutant) | *c.19G>A*<br>= E (Glu) > K (Lys) | AAG |
| HbSC (double mutant) | *HbS mutation (Glu→Val) +*<br>*HbC mutation (Glu→Lys)* | AAG + GTG [113] |

Abbreviations: Hb, hemoglobin ; HBB, human hemoglobin beta.

In samples collected at UZA, the HBB genotype sequences were determined by capillary hemoglobin electrophoresis at UZA as described elsewhere [112].

### *Ex vivo* sexual conversion assays

#### *Ex vivo* sexual conversion in *Plasmodium falciparum* field isolates

*P. falciparum*-infected fresh venous blood samples (15 ml) were collected from individuals carrying different HBB genotypes (Table 2; HbAA *n* = 14, HbAC *n* = 12, HbAS *n* = 6; total *n* = 32) and were cultured *ex vivo* to measure SCRs in natural human infections. Assay optimization and validation described in *S1 Appendix*. Cultures at 3% Hct and 20 ml volume were prepared by mixing 600 µl of *P. falciparum*-iRBC pellet with 20 ml of RPMI 1640 medium containing 0.5% Albumax II (Gibco, 11021-029) and 2 mM choline chloride (Sigma-Aldrich, C7527) [65, 81]. Parasites at the ring stage were incubated using candle jar method under static conditions until ∼24 hours post-culture (hpc). At this time, sexually committed rings present in the venous blood sample are expected to have developed into stage I gametocytes expressing Pfs16, while asexual parasites will have progressed to the schizont stage (Fig 2A). During culture, at ∼5 hpc, ML10 (80 nM), a cGMP-dependent protein kinase inhibitor, was added to prevent schizont rupture and reinvasion [45, 75]. Giemsa-stained smears were examined by LM to confirm that most parasites had reached either stage I gametocytes or schizonts stages. At ∼24 hpc, parasites were harvested and purified using MACS LD columns (Miltenyi Biotec). In brief, cultured parasites are centrifuged (1,648 g, 5 min) and pelleted iRBCs were diluted 1:4 with culture medium, loaded onto an LD-MACS column, and washed four times with 1 ml medium to separate non-iRBCs and ring-stage-iRBCs (first elution). The column was then removed from the magnet, and stage I gametocytes and schizonts were recovered by gravity elution, followed by gentle plunger flushing twice with 3 ml medium (second elution). Recovered parasites were pelleted (1,648 g, 5 min), resuspended 1:2 with non-iRBCs, and diluted 1:2 in PBS. From the second elution, four thin blood smears (3 µl each) were prepared and left to air-dry; three were used for on-site IFAs, and one was Giemsa-stained for LM to verify MACS enrichment.

#### Immunofluorescence assays

Immunofluorescence assays (IFAs) were performed at CRUN (Burkina Faso) as previously described [45, 65, 81], with modifications. Slides were stained in batches of four to eight cultures with three slides per culture, to optimize processing and staining consistency. Primary antibody mouse anti-Pfs16 (1:400, kindly provided by R. Sauerwein and T. Bousema, Radboud University) [76] and secondary antibody goat anti-mouse IgG (H+L) Alexa Fluor 488 (1:1000; Abcam, ab150113) were used, with incubations and washes under static conditions. Slides were mounted with Aqueous Fluoroshield (Abcam, ab104139) containing DAPI (0.0002%) and stored at 4°C until shipment to ITM (Belgium). At ITM, slides were visualized within six months of preparation on an ECHO Revolve epifluorescence microscope (ECHO, San Diego, CA) to determine SCRs (Fig 2B and 2C). Analysis was blinded to HBB genotype. Stage I gametocytes were identified as Pfs16-positive cells (green) and counted among a minimum of 1,000 DAPI-positive cells (blue) per slide (stage I gametocytes and schizonts). Slides with fewer than 1,000 DAPI-positive cells were excluded from further analysis during the first reading. For each culture (*n* = 35), three of the four prepared slides were stained by IFA and examined by a primary reader. Slides from three cultures were excluded due to insufficient DAPI-positive cells. From the remaining cultures (*n* = 32) a random subset of ∼10% (4/32), with three IFA slides per culture, was examined by a second reader to assess inter-reader reliability via intraclass correlation coefficient (ICC) analysis, based on a two-way random effects model, absolute agreement, single measures [ICC(2,1) [114]]. For each IFA slide, SCRs were calculated as the proportion of Pfs16-positive parasites among total DAPI-positive cells. Across the three replicate IFA slides per culture, the mean SCR (±SD) was computed and used to calculate and report the median SCR (IQR) by HBB genotype (Fig 2C), statistical differences were assess as described for proportions in section “Data analysis”. Fold-change in SCR for HbAC and HbAS was calculated relative to HbAA by dividing each mutant genotype’s median SCR by that of the wild-type. Statistical analysis was performed to assess differences in SCRs and in initial parasitemia across HBB genotypes (continuous not normally distributed variables) (Figs 2C-D), as well as the potential confounding effect of initial parasitemia on SCRs using non-parametric correlation (S1A Fig), as described in section “Data analysis”.

Stable Pfs16 fluorescence was confirmed by the counting of ≥1 stage I gametocyte per 10 fields. For each culture, the three replicate IFA slides were analyzed to quantify SCRs. The mean, minimum and maximum SCR were calculated across replicates. To assess inter-slide reproducibility (technical repeatability) of SCR measurements, an ICC (2,1) analysis (two-way random effects, single measures, consistency) was performed using SCR values from the three replicate slides per culture (*n* = 32). Counts of stage I gametocytes and total parasites were recorded for each IFA slide and summarized as median (Q1-Q3). From 5 out of 32 cultures (15.6%) specificity controls were performed by staining one slide per culture without primary antibody (negative controls). Stage V gametocytes (also express Pfs16), identified by their shape, were excluded from the counting as they are a product of SC occurring >10 days prior to the sexual commitment cycle under investigation. Giemsa-stained slides from each *ev*SCA were examined by LM to estimate parasite purification by MACS and confirm approximate parasite age/stage to assure parasites were sufficiently mature to express Pfs16 (starts expression at ∼20 hpi [45]).

### *In vitro* sexual conversion assays

*In vitro* SC assays (*iv*SCAs) were performed as follows (Fig 3A). NF54-*gexp02-Tom* parasites [65, 81] were routinely maintained with parasitemias of less than 2 % in O+ RBCs (3% Hct) in RPMI 1640-based medium supplemented with 0.5% Albumax II (Gibco, ref: 11021-029) and 2 mM choline chloride (+Choline; Sigma-Aldrich, C7527), analogous to LysoPC supplementation [60, 79, 81]. Cultures were incubated at 37°C with shaking conditions (70 rpm) in a gas mixture (5% O₂, 5.5% CO₂, balance N₂). To minimize bias due to prior culture duration, all cultures were initiated from parasites of the same cryopreserved batch of an expanded culture.

To obtain cultures of a well-defined age window, we used Percoll/sorbitol synchronization (Fig 3A). Percoll-purified schizonts (Generation −1) were used to establish fresh cultures with thawed cryopreserved RBCs (day 3 post thawing) with different HBB genotypes (exposure, Generation 0), now cultured in static conditions. Thawing of RBCs was performed using a three-step protocol, sequentially adding three volumes of 3.5% NaCl solution (10× the RBC volume each time), followed by RPMI washes and centrifugation (1,648 g, 5 min) as per standard procedure [115]. Approximately 5 hours later, when parasites have reinvaded the new RBCs, cultures were subjected to 5% D-sorbitol lysis [45, 81] to obtain cultures of a defined 0-5 hpi age window. Post-sorbitol initial parasitemia was then measured by flow cytometry, and parasites were cultured in two parallel subcultures (∼200 µl final volume) per sample. One sub-culture was maintained in +Choline medium (non-inducing SC conditions), and the other in choline-free medium (-Choline; inducing conditions). Cultures were maintained under these respective conditions for two generations (48 hpi, until 35-40 hpi in Generation 1, Fig 3A, hpi calculations are based on a 44 hours replication cycle empirically established for this parasite line in our lab). In each experiment we tested RBCs with at least two different HBB genotypes. Samples with different HBB genotypes were used in the experiments if the recovered RBC pellet volume after thawing was ≥ 30 µl after dilution with RPMI to 50% Hct. From a total of 20 cryopreserved non-iRBC samples from CRUN that were confirmed to be negative for *P. falciparum* by *var*ATS qPCR, 15 samples had a RBC pellet volume ≥ 30 µl after thawing and dilution in RPMI to 50% Hct (S7 Fig). The mean recovered RBC pellet volumes after thawing were: HbAA, 178 µl (range: 50−596 µl); HbAC, 111 µl (range: 30−320 µl); HbAS, 246 µl (range: 64−630 µl); HbSC, 558 µl (range: 35−1080 µl); and HbSS, 600 µl (range: 300−1120 µl). These samples were included within the *iv*SCAs, in addition to non-infected, cryopreserved RBC samples from the UZA HbSS and HbSC (S7 Fig). Total numbers of samples tested per HBB genotype and experiment are shown in S10 Table. Flow cytometry - analysis was used to measure parasitemia, assess parasite stage development (*i.e.,* proportion of parasites reaching the intraerythrocytic schizont stage or late-stage proportions), and quantify SC at defined time points during the assay (Fig 3A). Parasitemia was determined at 0-5 hpi (rings), together with (1) the proportion of schizont stages at 35-40 hpi in Generation 0 and 1, and (2) SCRs at 25-30 hpi (large rings/trophozoites) and 35-40 hpi (schizonts) in Generation 1 (Fig 3A). The standard timepoint to measure SC with this parasite line is 25-30 hpi to ensure detection of all sexually committed parasites, as TdTom-expression under the *gexp02* promoter starts around 10-15 hpi [81]. The 35-40 hpi timepoint was included at Generation 1 to account for potential delayed stage development in mutant HBB genotypes (see section on “Red blood cells with mutant HBB genotypes delay parasite development”). Four independent experiments were conducted, with RBC samples included in each experiment detailed in S10 Table. Each experiment included a fresh HbAA blood sample to control the effect of RBC storage after cryopreservation.

Samples for flow cytometry analysis (using a CytoFLEX LX flow cytometer, Beckman Coulter) were prepared by resuspending parasite cultures (1:3) in PBS containing 1% BSA, followed by staining with Hoechst 33342 (1:500, BD Pharmingen) in a final volume of 100 µl per well. Samples were then incubated at 37°C for 30 minutes. To measure parasitemia and proportion of schizont stages, flow cytometry analysis (with 100,000 events collected per sample) was set to detect Hoechst 33342 (350 nm laser; 450/40 filter). RBCs were gated by plotting side scatter area (SSC-A) versus forward scatter area (FSC-A), with singlets selected using FSC-H versus FSC-A. Parasites were identified within the singlet gate by Hoechst fluorescence. Hoechst-positive events were defined as exceeding background levels from unstained negative controls. To quantify SCRs, Hoechst fluorescence was combined with the fluorescent reporter TdTom (561 nm laser; 610/20 filter) that is integrated in the NF54-*gexp02-Tom* line to distinguish sexual (Hoechst+/tdTom+) from asexual (Hoechst+/tdTom-) parasites. SCRs were calculated as the proportion of Hoechst+/tdTom+ events among all Hoechst+ events.

SCRs were expressed as a percentage (median [IQR]) per HBB genotype (Figs 3B and S3A-C). Fold-changes in SCRs for HbAC, HbAS, HbSS and HbSC were calculated relative to HbAA by dividing median SCR of each mutant genotype group by that of the wild-type HbAA group. Statistical analysis was performed as described in the section on “Data analysis”, to assess differences in SCRs (Figs 3B and S3A-C) and potential confounding effect of post-sorbitol initial parasitemia (S1B Fig) across HBB genotypes. Reticulocyte percentages were measured for all RBC samples on the same day as Percoll synchronization (Generation −1, Fig 3A). Briefly, RBC suspensions were mixed 1:1 with reticulocyte stain (Sigma-Aldrich), incubated 15 min at room temperature, and used to prepare a thin blood smear for LM estimation of reticulocyte percentage. We determined the ABO and Rhesus D blood groups in all samples using agglutination tests with Seraclone™ Anti-A, Anti-B, and Anti-D (RH1) reagents (Bio-Rad).

### Parasite development in red blood cells with different HBB genotypes

Parasite stage development was evaluated during the *iv*SCAs (Fig 3A) at 35-40 hpi (Generation 0) and 25-30 hpi/35-40 hpi (Generation 1) using LM and flow cytometry. The proportion of rings, trophozoites and early/late schizonts was quantified at 35-40 hpi in Generation 0 and 25-30 hpi/35-40 hpi in Generation 1 by LM of Giemsa-stained thin blood smears (100 parasite counts per sample; Fig 4A) in a subset of RBC samples (HbAA *n* = 1; HbAC *n* = 1; HbAS *n* = 2; HbSS *n* = 2; HbSC *n* = 2). Flow cytometry was used as described earlier to determine the percentage of late stages. For this a distinct population of cells were gated with increased Hoechst 33342 fluorescence (increased DNA content due to multiplication of the nuclei in schizont stages), within the Hoechst+ cells (Fig 4B). Upon confirming consistent results between LM and flow cytometry after screening the same samples by flow cytometry, the same technique was subsequently used in all *iv*SCA samples (HbAA *n* = 5, HbAC *n* = 5 and HbAS *n* = 4, and HbSC *n* = 1 donors from a malaria-endemic area in Nanoro, Burkina Faso; HbSS *n* = 12, HbSC *n* = 1 and *n* = 6 control HbAA donors [HbAA CTRL] from a non-endemic area) to determine the proportion of late-stages (*i.e.,* schizonts) by HBB genotype (Figs 4B and 4C). From LM data we calculated: (1) the percentage of rings, trophozoites and early/late schizonts (Fig 4A), (2) the mean parasite age using the age in hpi [116] of each stage (rings, trophozoites, schizonts) and the observed proportions; and (3) the developmental delay (in hours) by subtracting mean parasite age of the mutant genotypes from the wild-type HbAA genotype. Finally, for each genotype, we compared the developmental delay between Generations 0 and 1 by calculating the difference in mean parasite age at 35-40 hpi between both generations. Then, the development of each independent NF54-*gexp02-Tom* culture was classified as: *expected development*, *delayed development* or *(partially) arrested/extremely delayed development* (see detailed analysis in *S1 Appendix)*.

### IgG and IgM detection by flow cytometry against *Plasmodium falciparum*

To study the IgG and IgM reactivity against *P. falciparum* in individuals with different HBB genotypes by flow cytometry, we selected plasma samples from humans living in a malaria-endemic area from the *InHost* longitudinal cohort study [74]. We classified samples as “endemic” (endemic plasma) or “non-endemic” (non-endemic plasma) for malaria based on the country of origin. An “endemic” country has consistent, stable *P. falciparum* transmission, while a “non-endemic” country has no local transmission, with infections typically imported [117]. As the expected variation and magnitude of potential differences among HBB genotypes were unknown, we based the sample size on the minimum of 8 individuals per HBB genotype used in the *ev*SCAs and *iv*SCAs. A maximum of 15-20 individuals per genotype was included to ensure sufficient sampling. Selection criteria included equal representation of the most frequent genotype groups (HbAA [*n* = 18], HbAC [*n* = 18], HbAS [*n* = 18], HbCC [*n* = 2], and HbSC [*n* = 5]), *P. falciparum* infection status with a large proportion of active *P. falciparum* infections (*n* = 10-15) and a smaller proportion of non-infected individuals (*n* = 2-5), age range (1-19 years to exclude maternal antibodies), and homogeneous distribution amongst the ethnicity (all Mossi) and village (Nanoro, Nazoanga, Soum, and Séguédin) groups. Table 2 shows the number of participants per HBB genotype included in the assays. The *P. falciparum* transgenic line NF54-*gexp02*-Tom [65, 81] was cultured under the same conditions as described in the *iv*SCAs section and Fig 3A, with the following modifications: (1) parasites were cultured only in wild-type HbAA fresh RBCs and (2) flow cytometry assays were performed on early trophozoite/stage I gametocyte-iRBCs at 20-25 hpi (Generation 1), following the protocol reported in Dantzler KW., et al., (2019) [82] with the adaptations described below.

Human plasma samples were pre-diluted in PBS with 1% Bovine Serum Albumin (BSA, Sigma Aldrich) and heparin (2,500U; Sigma Aldrich) (PBS-BSA-Hep) at dilutions of 1:5000 (IgG) and 1:100 (IgM). *In vitro* generated trophozoite- and stage I gametocyte-iRBCs (25-30hpi; Generation 1) were pre-diluted to 0.6% Hct in PBS-BSA-Hep. Pre-diluted human plasma samples were incubated at a 1:10 dilution with pre-diluted trophozoite- and stage I gametocyte-iRBCs, in separate 96-well plates for IgG and IgM assays for 1.5 hours at room temperature. After 2 washes with PBS-BSA-Hep, secondary antibodies were added to the trophozoite- and stage I gametocyte-iRBCs for 30 minutes at 4°C at 0.6% hematocrit. AlexaFluor488 (AF488) mouse anti-human IgG (Jackson ImmunoResearch Europe) and PerCP/Cy5.5® mouse anti-human IgM (Abcam) were used at final dilutions of 1:640 and 1:320 in PBS-BSA-Hep for the IgG and IgM assays, respectively. After 2 washes with PBS-BSA-Hep, the DNA stain Hoechst33342 (BD Pharmingen) was added to the trophozoite- and stage I gametocyte-iRBCs at a final dilution of 1:5000 in PBS-BSA-Hep at 0.6% hematocrit for 30 minutes at 37°C. To verify that the trophozoite- and stage I gametocyte-iRBCs measured contained exclusively RBCs (no white blood cells), each batch pre-diluted trophozoite- and stage I gametocyte-iRBCs (to 0.6% hematocrit in PBS-BSA-Hep) were stained with APC anti-human CD236 (Glycophorin C) Recombinant Antibody (Biolegend®) in triplicate at a 1:40 final dilution (S4C Fig). Immunostained samples were analyzed on the flow cytometer CytoFLEX LX (Beckman Coulter), collecting 100,000 cells per culture. We analyzed the median proportions (% cells) of total (trophozoites and stage I gametocytes), asexual (trophozoites) and sexual (stage I gametocytes) iRBCs that were positive for IgG and IgM binding to surface antigens on iRBCs.

Samples were considered positive for IgG or IgM if the total iRBC proportions exceeded that of a naïve non-endemic plasma from a volunteer (negative control with HbAA genotype with no prior *P. falciparum* infection, from a non-endemic country [*i.e.,* Belgium] collected at the clinic of the Institute of Tropical Medicine [ITM] in Antwerp). Three non-endemic samples from an acute *P. falciparum*-infected traveler collected at the ITM clinic at days 0, 21 and 23 of *P. falciparum* diagnosis (*i.e.,* acute *P. falciparum-infected* non-endemic plasma D0, D21 and D23) were used as positive controls. A schematic of the gating strategy to measure the surface reactivity of asexual (tdTom-) and sexual (tdTom+) iRBCs (Hoechst33342+) to IgG (IgG-positive trophozoite-iRBCs: Hoechst33342+/tdTom-/AF488+ and IgG-positive stage I gametocyte-iRBCs: Hoechst33342+/tdTom+/AF488+) or IgM (IgM-positive trophozoite-iRBCs: Hoechst33342+/tdTom-/PerCP-Cy5.5.+ and IgM-positive stage I gametocyte-iRBCs (Hoechst33342+/ tdTom+/PerCP-Cy5.5.+) in the analyzed endemic human plasma samples is represented in Figs S4B and S4C.

To determine the optimal plasma dilution factor, a pool of five endemic and hyperimmune plasma samples from *P. falciparum*-infected individuals in a sub-cohort within the *InHost* study [74] (*i.e., P. falciparum*-infected hyperimmune endemic plasma pool of symptomatic individuals with high *P. falciparum* parasitemia ranging from 45,667 to 175,000 parasites per µl) [66] was titrated for the IgG and IgM assay separately. For the IgM assay, four additional plasma samples were titrated (*i.e.,* “naïve non-endemic plasma” and “acute *P. falciparum*-infected non-endemic plasma at D0, D21 and D23”) (S4A Fig).

During flow cytometry analysis, the IgG- (Hoechst33342+/tdTom-/+/AF488+) and IgM-positive (Hoechst33342+/tdTom-/+/PerCP-Cy5.5.+) total iRBC proportions for each sample were expressed as total event counts, percentage of positive cells, and median fluorescence intensity (MFI) per HBB genotype group. The IgG- and IgM-positive iRBC proportions and signal were compared by HBB genotype group for the total (trophozoites and stage I gametocytes), asexual (trophozoites) and sexual (stage I gametocytes) iRBCs (Figs 5 and S5). To correct for the background signal from the non-iRBCs in the IgG and the IgM channel, the MFI of all samples (including the positive and the negative controls) was expressed as a ΔMFI. For the asexual and sexual iRBCs, ΔMFI was calculated as ΔMFI (AF488+/PerCP-Cy5.5.+) = (“MFI IgG+/IgM+ of asexual [tdTom-]/sexual [tdTom+] trophozoite-iRBCs [Hoechst33342+]”) - (“MFI IgG+/IgM+ of non-iRBCs [tdTom-/Hoechst 33342-]”) within each plasma sample. For the total iRBCs, ΔMFI was calculated as ΔMFI (AF488+/PerCP-Cy5.5.+) = (“MFI IgG+ or IgM+ iRBCs [Hoechst33342+]”) - (“MFI IgG+ or IgM+ of non-iRBCs [Hoechst33342-]” of trophozoite stages incubated with PBS-BSA-Hep, instead of plasma).

### Data analysis

All flow cytometry data were analyzed using FlowLogic™ version 8.7 (Inivai Technologies™). Statistical analyses were performed in R version >4.5.1 (R Core Team 2025), RStudio software version 2025.09.1+401(©2025 Posit Software; RRID:SCR_000432) and the following R packages: brms v. 2.23.0 [118–120], DHARMa v. 0.4.7 [121], emmeans v. 1.11.2.8 [122],ggplot2 v.4.00 [123], glmmTMB v. 1.1.13 [124, 125], here v. 1.0.2 [126], lme4 v. 1.1.37 [127], performance v. 0.15.2 [128], renv v. 1.1.5 [129], rmarkdown v. 2.30 [130–132], scales v. 1.4.0 [133], sjPlot v. 2.9.0 [134], tidyverse v. 2.0.0 [135], with p-values (*p* < 0.05) considered significant.

Risk analysis was conducted to examine associations of demographic and clinical variables (*e.g.,* gender, age, village, ethnicity) across HBB genotype groups (S2, S4 and S8 Tables for the *ev*SCAs, *iv*SCAs, and IgG and IgM detection, respectively). An additional risk analysis for factors associated with elevated SCRs included the same covariates with the addition of HBB genotype (Table 1, for the *iv*SCAs). Continuous variables were analyzed using the Kruskal-Wallis test when parametric assumptions were not met, and when significant, this was followed by Wilcoxon rank sum post-hoc tests for pairwise comparisons with Holm correction. Otherwise, One-way ANOVA or Welch’s ANOVA was applied depending on whether Bartlett’s test indicated equal or unequal variances, respectively. When significant, Bonferroni-corrected post-hoc pairwise comparisons were performed. Associations between categorical variables were evaluated using Fisher’s exact test. Spearman correlation was used for correlation between continuous variables with no normal distribution.

For modelling SCR, parasite stage proportions, initial parasitemia and reticulocyte proportions in *ev*SCAs, and *iv*SCAs, and IgG/IgM proportions in the flow cytometry assays, the statistical analysis type was chosen based on the nature of the variables: proportions based on counts were assessed using beta-binomial generalized linear models (GLM) and mixed models (GLMM) with Bonferroni adjustment for the odds ratio contrasts in pairwise comparisons. We chose beta-binomial models due to (possible) violations of the assumptions of the binomial model (namely overdispersion). For *ev*SCA, random factors were used to account for technical replicates and a per-HBB-genotype dispersion factor was added to the GLMM model to account for heteroskedasticity. To address the liberal parametric Wald Z estimates and confidence intervals (asymptotic approximations) provided by the Generalized linear (mixed) models [GL(M)M models], we also computed more conservative Bayesian estimates, odds ratios and credible intervals using a Markov chain Monte Carlo (MCMC) approach when model specification and convergence allowed. GLM and GLMM model specifications and analysis results are reported in S1A-C Table. When the assumptions of the parametric beta-binomial models were violated (*e.g.,* dispersion or quantile deviations as assessed by DHARMa), we instead performed non-parametric Kruskal-Wallis and pairwise Wilcoxon rank-sum tests as described below.

Variable selection was guided by a focus on the primary biological variable of interest while avoiding model overspecification and multicollinearity, given the small sample size and unbalanced group structure in terms of potential confounding factors. Our primary analyses therefore focused on the association between SCR, HBB genotype, parasite stages, parasitemia and reticulocyte proportions.

For each analysis, we considered additional demographic and clinical variables as potential confounders. Univariate risk analysis of the *iv*SCA data suggested possible associations between SCR and other variables (Table S4). Originally correcting for confounding factors was not part of the experimental design and sample size calculations which resulted in low number of observations per level and sparse and uneven representation in the *iv*SCA. Consequently we were unable to properly control for those variables in the GL(M)M analyses (or without violating modelling assumptions in the case of *ev*SCA), Moreover, likelihood ratio tests did not indicate improved model fit when additional variables were included, compared to models using HBB genotype as the sole independent variable. We therefore focused on a descriptive modelling approach and prioritized parsimonious models with a single parameter of interest, which provided the most interpretable inference given the constraints posed by the available data.

Although this approach cannot fully exclude confounding or interaction effects, the direction and magnitude of estimated effects were broadly consistent across alternative model specifications. Sample size and power limitations should still be kept in mind when interpreting some of these GL(M)M results, with emphasis on effect sizes over exact p-values.

### Ethical approvals

The *InHost* study, a longitudinal cohort in Burkina Faso, was approved by the ethical review board of the Comité d’Éthique pour la Recherche en Santé (CERS: 2018/10/131) in Burkina Faso, and by the IRB-ITM (1261/18) and IRB-UZA (19/06/064) in Belgium. The amendment for our nested study in the *InHost* study was approved by CERS (N°2022000140/MSHP/ MESRI/CERS), the IRB at ITM (1261/18), and the IRB at UZA (3347) in Belgium. Ethical approval for additional sample collection at the Hematology Unit at UZA was granted by the IRBs at ITM (1538/21) and UZA (3215).

## ACKNOWLEDGEMENTS

We are grateful to Robert W Sauerwein and Teun Bousema (Radboud University, The Netherlands) for the anti-Pfs16 monoclonal antibody. We thank Núria Casas-Vila (ISGlobal) for the support and guidance in setting up the *ex vivo* and *in vitro* SC assays. We thank Eduard Rovira-Vallbona (ISGlobal) for helpful discussions on hemoglobin beta variants and aspects of the *in vitro* study.

## SUPPORTING INFORMATION

**S1 Appendix:** Assay optimization and validation of *ex vivo* and *in vitro* SC assays, and IgG/IgM detection by flow cytometry.

**S1 Fig. Correlation analysis between parasitemia and SC rates (SCRs).** (A) *ev*SCAs and (B) *iv*SCAs. Data points are colored by HBB genotype. Spearman’s rank correlation was used; non-significant correlations are indicated as ns.

**S2 Fig.** Representative flow cytometry plots showing tdTomato and Hoechst33342 signals. (A) HbAA fresh blood controls cultured under +Choline and -Choline conditions, with and without Hoechst33342. (B-C) Parasites cultured in cryopreserved RBCs with different HBB genotypes under +Choline (B) or -Choline (C) conditions.

**S3 Fig. Effect of HBB genotype on SC in *in vitro* SC assays (*iv*SCAs).** Related to Fig 3. SCRs were measured by flow cytometry at 25-30 hpi in +Choline and -Choline cultures (A-B) and at 35-40 hpi in -Choline cultures (C). (D-E) Show the proportions of asexual versus sexual stages and the correlation between reticulocyte percentage and SCRs. Statistical tests: (B-C) beta-binomial GLMs with Bonferroni correction; (A) Kruskal-Wallis test; (E) Spearman correlation.

**S4 Fig.** Flow cytometry detection of IgG and IgM responses against *P. falciparum*-iRBCs. (A) Plasma titration curves with positive and negative controls. (B-C) Gating strategy for measuring antibody reactivity to asexual (tdTom-) and sexual (tdTom+) iRBCs and verification of RBC-only populations using Glycophorin C.

**S5 Fig. IgG- and IgM-mediated responses to *Plasmodium falciparum* trophozoites and stage I gametocytes across HBB genotypes.** (A-D) Event counts and MFI for IgG- and IgM-positive total, asexual, and sexual iRBCs in plasma from *P. falciparum*-infected and non-infected individuals. Statistical comparisons were performed with beta-binomial GLMs with Bonferroni correction.

**S6 Fig. IgG- and IgM-positive total iRBCs in plasma from *P. falciparum*-infected and non-infected individuals.** Median proportions of IgG-positive (A) or IgM-positive (B) total-iRBCs for *Pf*-infected versus non-infected groups. Statistical comparisons were performed with beta-binomial GLMs with Bonferroni correction.

**S7 Fig. Flow chart of participant selection in the *ex vivo* and *in vitro* SC assays.** Infection status determined by RDT, LM, and *var*ATS qPCR. Samples meeting parasitemia and RBC volume criteria were included in respective assays.

**S8 Fig. Experimental variability of SCR measurements in +Choline cultures, grouped by HBB genotype.** Timepoint of measurement: at 35-40 hpi (Generation 1).

**S1A Table.** Statistical analysis outcomes from GLMs or GLMMs and Kruskal-Wallis tests in *ex vivo* SC assays.

**S1B Table.** Statistical analysis outcomes from GLMs or GLMMs and Kruskal-Wallis test in the *in vitro* SC assays.

**S1C Table. Statistical analysis outcomes from GLMs or GLMMs and Kruskal-Wallis tests in the IgG/IgM detection assays.**

**S2 Table. Risk analysis of SCRs and demographic/clinical variables across HBB genotypes in *ev*SCAs.** Data from *P. falciparum*-infected samples (parasitemia ≥ 0.05% or 2500 parasites/µl) collected in the Nanoro Department, Burkina Faso. Statistical tests: Chi Square /Fisher exact test, Kruskal Wallis test.

**S3 Table.** Median SCRs (IQR) by HBB genotype measured in *iv*SCAs with flow cytometry. Timepoints measured: at 25-30 hpi and 35-40 hpi (Generation 1).

**S4 Table.** Risk analysis of factors across HBB genotype groups, in *iv*SCAs.

**S5 Table. RBC counts per µl analyzed during flow cytometry to exclude rule hemolysis of fragile mutant RBCs.** Measurements were done at three timepoints: 35-40 hpi (Generation 0), and 25-30 hpi and 35-40 hpi (Generation1), for HbAC; HbAS, HbSS and HbSC RBCs. Statistical test: Kruskal-Wallis.

**S6 Table. Analysis of parasite stage development by individual parasite culture within each HBB genotype group.** Parameters (PMR, late-stage proportions, fold changes) compared to HbAA reference and color-coded. Cultures classified as expected, delayed, or (partially) arrested/extremely delayed. SCR at 35-40 hpi (Generation 1) also shown.

**S7 Table. Median IgG- and IgM-positive proportions of total, asexual, and sexual iRBCs by HBB genotype.** Measurements done by flow cytometry.

**S8 Table. Demographics of participants in IgG/IgM detection assays across HBB genotype groups.** Data was analyzed in *P. falciparum* infected and non-infected plasma samples collected in the Nanoro Department, Burkina Faso within the context of the *InHost* study.

**S9 Table.** Key resources, primers, and probe sequences.

**S10 Table.** Overview of experiments and number of cryopreserved RBC samples by HBB genotype.

