## Supporting information for "Elevated *Plasmodium falciparum* sexual conversion in HbAC and HbAS red blood cells in naturally infected malaria patients"

### S1 Appendix: Assay optimization and validation

#### *Ex vivo* sexual conversion assays

To ensure robustness and field feasibility of the *ex vivo* sexual conversion assays (*ev*SCAs) using *P. falciparum* isolates with varying parasitemias, we optimized key parameters using mock cultures of the NF54-*gexp02-Tom* transgenic line [1] (Fig 2A). Method optimization focused on: 1) parasite purification and enrichment methods suitable for field conditions, and 2) assessment of the minimal detectable parasitemia and culture volume for accurate SCR quantification.

First, for parasite purification we compared the magnetic-activated cell separation (MACS) LD and LS columns (Miltenyi Biotec) using mixed stages of NF54-*gexp02-Tom* parasites (10 ml culture volume, 3% hematocrit [Hct], ~11% parasitemia). In brief, ~255-300 µl of infected-RBC (iRBC) pellet was diluted 1:4 with culture medium (final volume: ~1,020-1,200 µl) to reduce viscosity and facilitate efficient separation, and passed through the columns. We performed two elution steps with culture medium: the first elution (6 ml) containing non-infected RBCs and ring stages; and a second elution (6 ml) with the bound fraction after removing the column from the magnet yielding hemozoin-containing stage I gametocytes and schizonts. The first and second elution were then pelleted (1,648 g, 5 min centrifugation). From the purified RBC pellets, 3 µl (LD column) and 1.5 µl (LS column) were used to prepare thin blood smears to quantify the percentage of parasite stages by light microscopy (LM), and to confirm the presence of ring stages (first elution) and gametocytes stage I and schizonts (second elution). Results showed better purification with LD columns, yielding a larger RBC pellet (~8 µl vs. ~1.5 µl) and higher parasite enrichment (~75% vs. ~58%). LD columns were therefore used for all subsequent steps.

Second, to determine the minimal detectable parasitemia, NF54-*gexp02-Tom* parasites were cultured at 3% Hct in 10 and 20 ml volumes to a parasitemia of ~1% iRBC, followed by a 10-fold serial dilution with non-infected RBCs to obtain parasitemias from ~1% to ~0.001%. Then, for each condition (dilution point) RBC pellets were diluted 1:4 with culture medium (end volume of ~1,020 µl) and purified with MACS LD columns. A thin blood smear per condition was prepared from 3 µl of the purified RBC pellet without dilution, followed by IFAs to quantify Pfs16-positive parasites. For fresh (*i.e.,* non-cryopreserved) infected red blood cells (iRBCs), the minimum detectable parasitemia was 0.1% in 10 ml cultures (~3 million parasites) and 0.01% in 20 ml cultures (~600,000 parasites).

Additionally, we compared the minimal detectable parasitemia between matched cultures with fresh and cryopreserved iRBCs at 1% parasitemia (3% Hct, 20 ml culture, and 600 µl RBC pellet at 100 % Hct) to evaluate whether sexual conversion assays (SCAs) could be performed without loss of sensitivity after parasite cryopreservation. Cryopreservation and thawing was done as described in section below *"In vitro sexual conversion assays": RBC integrity after thawing across human hemoglobin beta (HBB) genotypes"*. In cryopreserved iRBCs, matched to cultures with non-cryopreserved iRBCs (same total volume, RBCs pellet volume and number of parasites before cryopreservation), the detection threshold increased to 0.1% (~6 million parasites), due to reduced parasitemia after cryopreservation (1 % to ~0.17%). These findings identified 20 ml fresh iRBC cultures as the optimal setup for subsequent assays.

Lastly, to ensure comparable sexual conversion (SC) measurements across smears regardless of parasitemia, we optimized conditions to obtain a homogeneous monolayer of infected red blood cells (iRBCs) mixed with non-iRBCs in the thin blood smears prepared after MACS. We evaluated different dilutions of the MACS-purified iRBC pellet in non-iRBCs and PBS using fresh mock cultures (3% hematocrit, 20 ml) with 10-fold serial parasitemia dilutions (~1% to ~0.01%),. The most effective strategy resulted in a two-step dilution: first 1:2 with fresh non-iRBCs, then 1:2 with 1× PBS. The final *ev*SCA protocol was validated on clinical *P. falciparum* samples cultured *ex vivo* (3% Hct, 20 ml fresh culture) at the Clinical research unit in Nanoro, Burkina Faso (CRUN), confirming field feasibility and reliable detection of sexual parasites at parasitemias as low as 0.05% (vs. 0.01% in mock samples). Among 32 successful *ev*SCAs, 41% (13/32) had starting parasitemias of 0.05-0.08%, and the remaining 59% (19/32) ranged from 0.08-1%.

Inter-reader reliability was assessed by re-reading three IFA slides per culture in 12.5% of the total cultures (4/32) by a second reader. Intraclass correlation coefficient (ICC) analysis of the resulting SC rates (SCRs), demonstrated very strong inter-reader agreement with an ICC (2,1) (two-way random effects, single measures, consistency) of 0.96 (95% CI: 0.543-0.998, $p$ = 0.0055**). IFA smears showed consistent Pfs16 fluorescence, with a median of 44 stage I gametocytes per smear (IQR: 28-64), an average of 19 stage I gametocytes per 10 fields (confirming at least one visible stage I gametocyte per 10 fields), a median of 1,021 total parasites counted per smear (IQR: 30, with Q1: 1,009 and Q3: 1,039), and 20 counted fields per smear (range: 14-32). Inter-smear reproducibility of SCRs across three IFA replicate slides per culture showed good consistency (ICC of 0.79 [95% CI: 0.66-0.88]; $p < 0.001$). Giemsa-stained replicate smears confirmed a MACS enrichment efficiency of ~70% and that ~54% of parasites observed after MACS were late schizonts [2]. Therefore, if circulating asexual rings had sufficient *ex vivo* culture time to progress to the schizont stage, it can be assumed that circulating sexual rings had sufficient time to progress to Pfs16-expressing stage I gametocytes (note that progression to stage I gametocytes takes a similar time to progression to trophozoites).

#### *In vitro* sexual conversion assays

##### Red blood cell integrity after thawing across HBB genotypes

To validate the experimental set-up of the *in vitro* SC assay (*iv*SCA), we first evaluated RBC integrity and recovery after thawing across a subset of RBC samples with different genotypes

($n$ = 1 for each cryopreserved RBC sample from a malaria-endemic area in Nanoro, Burkina Faso (CRUN): hemoglobin AA [HbAA], HbAC, and HbAS, compared to fresh HbAA RBCs of donors from a non-endemic area at the Institute of Tropical medicine in Antwerp, Belgium (ITM). Thawing was performed using a three-step protocol, sequentially adding three volumes of 3.5% NaCl solution (10× the RBC volume each time), followed by RPMI washes and centrifugation (1,648 g, 5 min) as per standard procedure [3]. Since RBCs with mutant HBB genotypes are more fragile and lysis-prone, hemolysis was assessed across HBB genotypes on day 0 to evaluate whether the thawing procedure led to RBC lysis and at days 0, 3 and 7 to assess for how long the RBCs could be used for culture after thawing. The recovered RBC pellet volume (µl) at day 0 (after thawing) was measured using a pipette and the number of RBCs per milliliter was assessed using the hemocytometer [Scepter™ 3.0 Handheld Cell Counter, Millipore® Cat: PHCC30000]) on day 0, 3, and 7 (after thawing). The minimum required volume of recovered RBC pellet at day 0 (after thawing) to initiate culture wells was at least 30 µl after dilution with RPMI to 50% Hct, samples below this threshold were removed (S7 Fig). RBC counts per milliliter at day 0, 3 and 7 were: fresh HbAA RBCs (non-endemic donors), 3.61e+09, 9.65e+09 and 3.35e+09; cryopreserved HbAA RBCs, 1.06e+10, 1.73e+09 and 3.31e+09; cryopreserved HbAC RBCs, 4.62e+09; 5.41e+09 and 2.67e+09; and cryopreserved HbAS RBCs, 7.07e+10, 1.10e+10 and 3.35e+09. RBC counts decreased by day 7 across all HBB genotypes and thawed RBCs at day 0 appeared morphologically distressed by LM right after the thawing procedure; therefore, we used RBCs three days after thawing for the *iv*SCAs.

Finally, as the cryopreserved RBC samples collected at CRUN and the UZA were stored in differently (CRUN: at -80°C before transfer to liquid nitrogen at ITM, three months after sample collection; UZA: directly snap-frozen in liquid nitrogen at ITM) we assessed if this difference in storage had an impact on RBC lysis by comparison of the recovered RBC pellet volume (µl) in three donor-matched (non-endemic) HbAA control samples, preserved according to either the CRUN or UZA protocol, or used fresh. No differences were found in the two cryopreserved samples as these yielded 200 µl of recovered RBC pellet (at 100% Hct) after thawing from an input cryopreserved volume of RBCs at 100% Hct of 500µl (40% of RBC pellet recovery after cryopreservation), compared to 1,000 µl of RBCs at 100% Hct obtained in fresh RBCs. Importantly, SCRs were similar across conditions: 8.5% in fresh HbAA RBCs, and 7.7% versus 6.8% in the cryopreserved HbAA RBCs with storage according to CRUN and UZA, respectively. The results suggest that neither cryopreservation nor subsequent storage methods affected RBC integrity after thawing or SCRs in the *iv*SCA. SCRs showed comparable variability within HBB genotypes across experiments, with tightly clustered values in wild-type HbAA controls (median 8.11 and IQR 1.36; S8 Fig)

#### Parasite development in red blood cells of different HBB genotypes

Parasite development in RBCs with different HBB genotypes was assessed during the *iv*SCAs using LM and flow cytometry at multiple time points (25-30 hpi and 35-40 hpi) across two asexual replication cycles (Generation 0 and 1; Fig 3A). In Generation 0 (35-40 hpi, Fig 4A left graph), parasites in wild-type HbAA RBCs reached the expected mean parasite age of ~43 hpi, closely matching the ~44 hpi life cycle, with parasite stage distributions dominated by schizonts (87%). In contrast, mutant HBB RBCs showed genotype-specific developmental delays with mean parasite ages of ~40 hpi (HbAC), ~33 hpi (HbAS), ~34 hpi (HbSS), and ~28 hpi (HbSC), corresponding to developmental delays of ~3-15 hours. These delays were reflected in parasite stage distributions, with a marked shift toward earlier stages (increased proportions of trophozoites and rings relative to HbAA), most pronounced in HbSC. Similar delays persisted in Generation 1 (25-30 hpi [first timepoint of SCR measurement] and ~35-40 hpi; Fig 4A middle and right graph), with mean parasite ages further reduced compared to Generation 0, indicating continued developmental delays and prolonged life cycles. Even in HbAA, mean parasite age was ~38 hpi (*i.e.,* ~6 hours younger than the expected ~44 hpi cycle), with parasite stage proportions shifting toward earlier stages (60% schizonts versus 87% in Generation 0). Developmental delays relative to HbAA persisted in mutant genotypes: ~2 hours in HbAC, ~9 in HbAS, ~7 in HbSS, and ~13 in HbSC, again supported by parasite stage distributions showing a shift towards earlier stages (rings/trophozoites) and reduced schizont proportions. Notably, a subset of schizont stages, potentially persisting from Generation 0 in HbSS and HbSC RBCs, were observed across timepoints suggesting that a subset of parasites may remain developmentally arrested, besides delayed development.

Flow cytometry analysis of late stage (schizonts) proportions at 35-40 hpi confirmed LM observations. In Generation 1 at 35-40 hpi, late stage proportions were significantly lower in HbSS compared to HbAA RBCs (beta-binomial GLM, OR[95% CI] = 0.13 [0.03−0.60], *p [Bonferroni]* = 0.0026; Fig 4B, right graph and S1B Table), while HbAS and HbSC RBCs showed non-significant reductions (Beta-binomial GLM; $p = 1$; Fig 4B, right graph). Same trends were observed in Generation 0 at 35-40 hpi, but were not-significant (Kruskal-Wallis test, overall $p = 0.05$; Fig 4B, right graph). These findings support our earlier LM-based observations, pointing toward a slight developmental delay in parasites within HbAS and HbSC RBCs not prone to impact SCR measurements with the NF54-*gexp02-Tom* line and a significant delay in HbSS RBCs that could impact SCR measurements. At 25-30 hpi (Generation 1), parasites in all HBB genotypes were predominantly at early stages (rings/trophozoites), as expected, indicated by low late stage proportions (Fig 4B, middle graph). At this timepoint, no significant differences were observed between genotypes (Kruskal Wallis test, $p = 0.13$). The progression from predominantly late stages at 35-40 hpi in Generation 0 to early stages at 25-30 hpi (Generation 1) to late stages in at 35-40 hpi in Generation 1 shows that in HbAA, HbAC and HbAS RBCs there was no complete arrest of parasite development. Notably, in mutant

HbSS and HbSC RBCs at 25-30 hpi (Generation 1), a proportion of late stages was detected by LM that is absent in other HBB genotypes (Fig 4A, middle graph), yet this population is not detected by flow cytometry (Fig 4B, middle graph). This may suggest that the observed schizonts might be underdeveloped (potentially containing less DNA and thus not being detected by flow cytometry as late stages).Parasite multiplication rates (PMR) analyzed by HBB genotype confirmed parasite development and multiplication across Generation 0 to 1, with median [IQR] PMRs >1 for all genotypes and slightly lower values in HBB mutant genotypes compared to HbAA (HbAA, 3.5 [2.0]; HbAC, 2.0 [1.0]; HbAS, 3.5 [1.5]; HbSS, 4.0 [2.0]; and HbSC, 2.0 [1.0]) (Kruskal-Wallis test, $p = 0.83$).

As the proportions of late stages showed high IQRs per genotype at the different time points, we further investigated whether delay or arrest was more pronounced in certain cultures, taking into account that other host factors may vary from participant donating the RBCs apart from HBB genotype. We analyzed several parameters obtained by flow cytometry: the proportion of late stages at three timepoints (35-40 hpi at Generation 0; 25-30 hpi at Generation 1; and 35-40 hpi at Generation 1), and fold change of late stages from one time point to the next, as well as the PMR (S6 Table). We defined three developmental classes based on the comparison of the median and interquartile ranges of each parameter in parasites growing in each group of mutant HBB RBCs compared to the HbAA RBCs: *expected development* (≤ one parameter outside the HbAA reference range), *delayed development* (2-3 parameters outside the HbAA reference range); *(partially) arrested/extremely delayed* development (≥ 4 parameters outside the HbAA reference range). In HbAA (5/5 [100%]) and HbAA CTRL RBCs (6/6 [100%]), all cultures exhibited the expected developmental pattern. In contrast, the HBB mutant RBCs showed variable developmental patterns: HbAC −40% (2/5) expected, 60% (3/5) delayed; HbAS−50% (2/4) expected, 25% (1/4) delayed, 25% (1/4) (partial) arrest/extremely delayed; HbSS −0%

(0/12) expected, 33% (4/2) delayed, 67% (8/12) (partial) arrest/extremely delayed; HbSC −0% (0/2) expected, 50% (1/2) delayed, 50% (1/2) (partial) arrest/extremely delayed (S6 Table).

#### IgG and IgM detection by flow cytometry

To establish the assay for IgG and IgM detection by flow cytometry, plasma titration was performed. Plasma samples from individuals living in malaria-endemic area were defined as "endemic plasma", whereas samples from individuals living in non-endemic area were defined as "non-endemic plasma". The plasma titration curves showed a clear dose-response dynamic for IgG detection with the "*P. falciparum*-infected hyperimmune endemic plasma pool", while for IgM detection the measured percentage of positive cells was low even at the lowest plasma dilution. Therefore, four additional plasma samples were titrated for IgM detection (*i.e.,* "naïve non-endemic plasma" and "acute *P. falciparum-infected* non-endemic plasma at D0, D21 and D23"). While the "naïve non-endemic plasma" remained under the threshold of the "hyperimmune *P. falciparum*-infected endemic plasma pool", the "acute *P. falciparum*-infected non-endemic plasma at D21 and D23" exceeded the values of both the "*P. falciparum*-infected hyperimmune endemic plasma pool" and the "naïve non-endemic plasma" and showed a dose-response dynamic. Due to the generally low proportions of IgM-positive early trophozoite/stage I gametocyte-iRBCs, we chose the optimal plasma dilution with highest percentage of positive cells detected for IgM (1:320), while for IgG detection we used the dilution in the slope (1:640) (S4 Fig).

### FIGURES AND TABLES

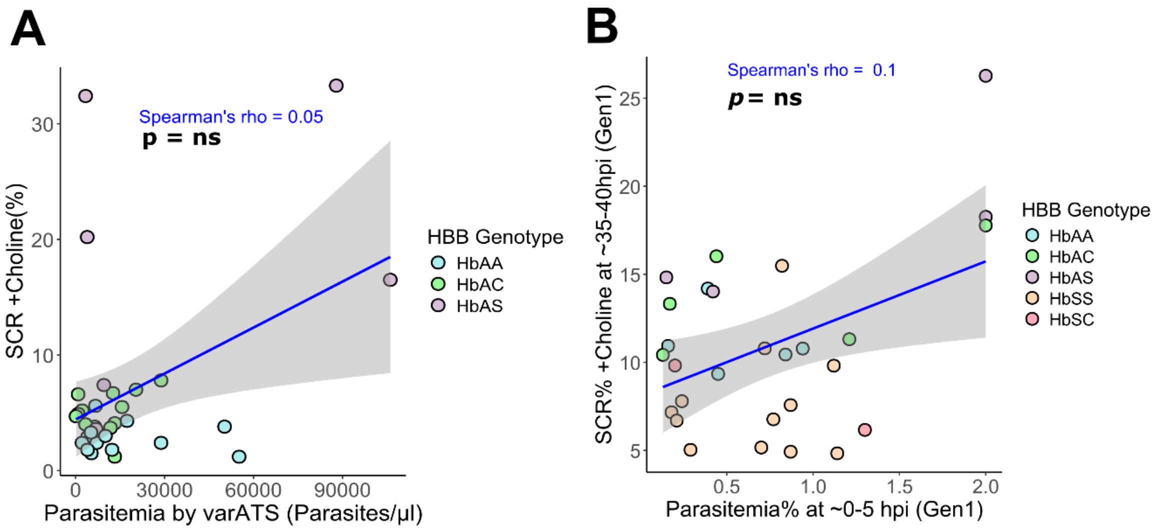


**S1 Fig. Correlation analysis between parasitemia and sexual conversion rates. (A)** Correlation between parasitemia at the start of the *ex vivo* sexual conversion assays (*ev*SCAs), determined by *var*ATS qPCR, and SCRs. **(B)** Correlation between parasitemia at the start of the *in vitro* sexual conversion assays (*iv*SCAs), at ring stages 0-5 hpi (Generation 1), and SCRs measured at 35-40 hours post-invasion (hpi; Generation 1). Data points are colored by HBB genotype. Statistical correlations were assessed using Spearman's rank correlation. Non-significant correlations are indicated as "ns". **Abbreviations**: Gen, generation; Hb, hemoglobin; HBB, human hemoglobin beta; hpi, hours post-invasion; *var*ATS qPCR, *var* gene acidic terminal sequence (*var*ATS) quantitative PCR (qPCR).

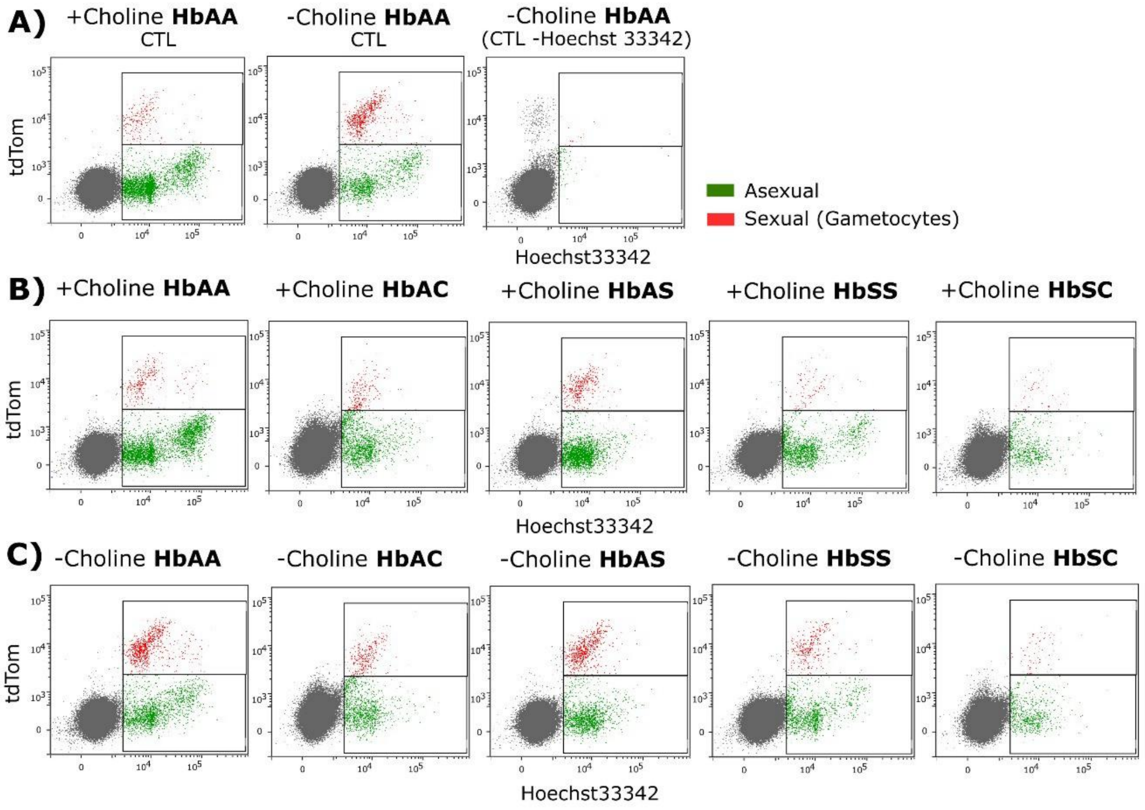


**S2 Fig. Representative tdTom (marks sexual stages/gametocytes) versus Hoechst33342 (stains parasite DNA) flow cytometry plots. (A)** Parasites in HbAA fresh blood control (CTL) cultured in +Choline and -Choline conditions both marked with the DNA stain Hoechst33342 (left and middle plots, respectively); and -Choline with no addition of the DNA stain Hoechst33342 (right plot). **(B)** Parasites cultured in cryopreserved RBCs with different HBB genotypes in +Choline culture medium, and **(C)** in -Choline culture medium (same samples as in panel B, parallel cultures). **Abbreviations:** CTL, control; Hb, hemoglobin; HBB, human hemoglobin beta; RBCs, red blood cells; tdTom, TdTomato.

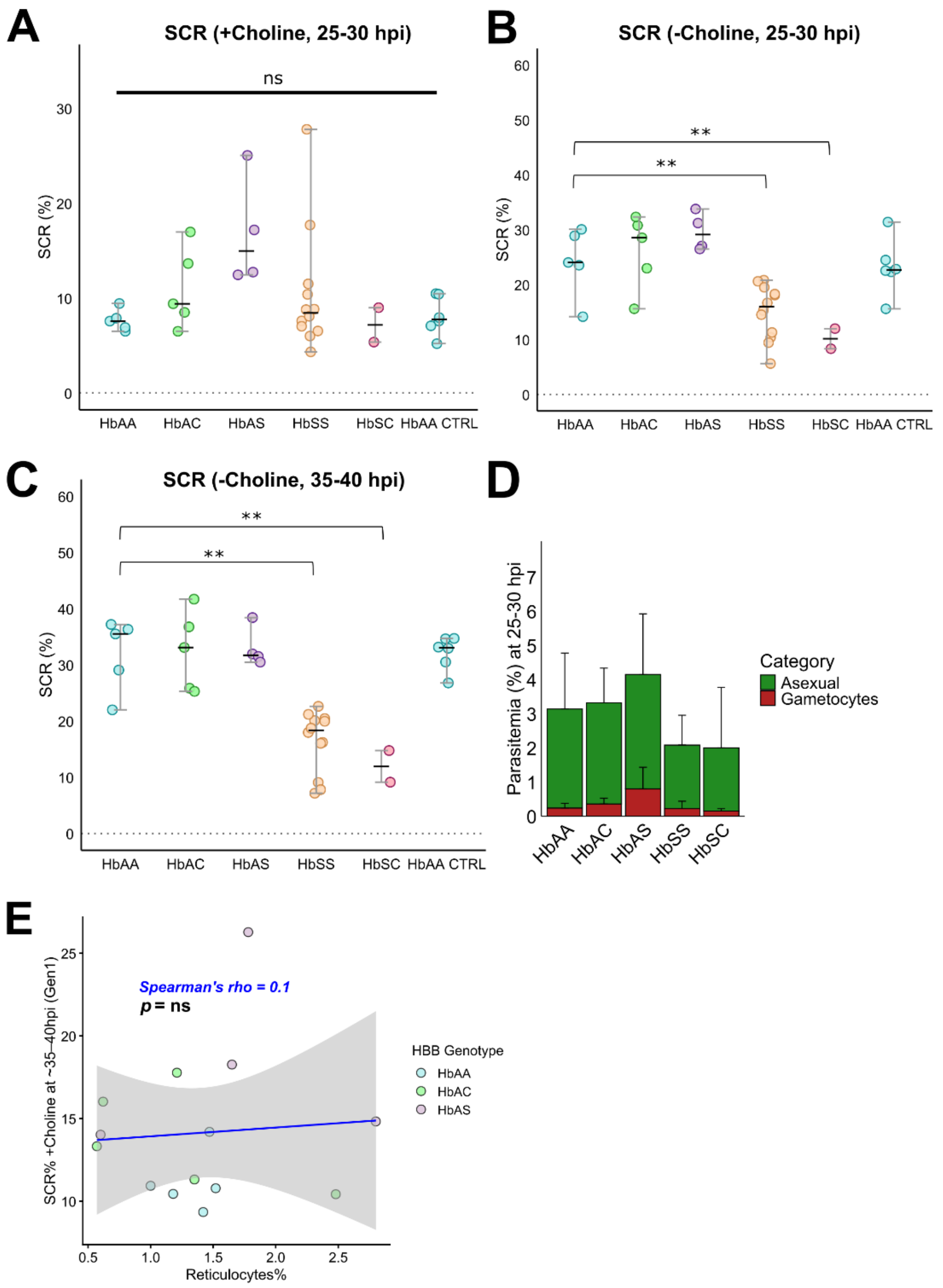


**S3 Fig. Effect of HBB genotype on SC in *in vitro* SC assays (*iv*SCAs). (A-E)** Sample size *n* per group: HbAA *n* = 5, HbAC *n* = 5 and HbAS *n* = 4 (endemic areas); and HbSS *n* = 12,HbSC *n* = 2 and "HbAA CTRL" *n* = 6 (non-endemic areas). **(F)** Sample size *n* per group: HbAA *n* = 5, HbAC *n* = 5 and HbAS *n* = 4 (endemic area). Sexual conversion rates (SCRs) by HBB genotype determined with flow cytometry at **(A)** 25-30 hpi (Generation 1) in +Choline cultures; **(B)** 25-30 hpi (Generation 1) in -Choline cultures; and **(C)** 35-40 hpi (Generation 1) in -Choline cultures. **(D)** Proportion of asexual and gametocyte stages within total parasitemia by HBB genotype at 25-30 hpi (Generation 1), in +Choline cultures. **(E)** Correlation analysis between reticulocytes percentage (X-axis) and SCRs at 35-40 hpi in +Choline cultures. Data points are colored by HBB genotype. **Statistical tests:** B-C, beta-binomial GLMs with Bonferroni adjustment for pairwise comparisons; (A) Kruskal-Wallis test; (E) Spearman correlation. Significant p-values are shown as: $p < 0.05$*, $p < 0.01$**, $p < 0.001$***. **Abbreviations:** CTRL, control sample (fresh HbAA RBCs); GLMs, generalized linear models; Hb, hemoglobin; HBB, human hemoglobin beta; Hpi, hours post-invasion; SC, sexual conversion; SCRs, sexual conversion rates.

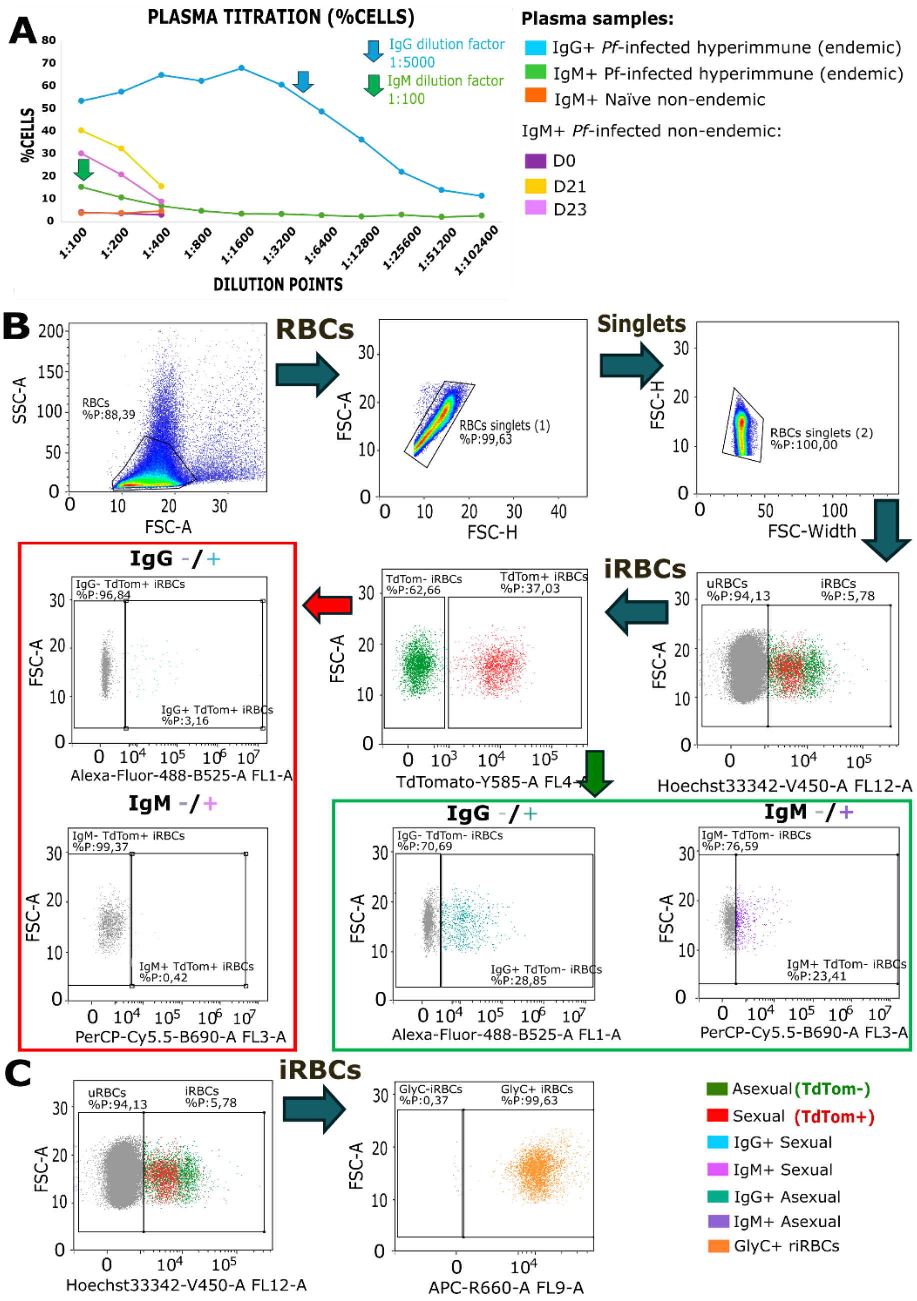


**S4 Fig. Flow cytometry detection of IgG and IgM responses against *P. falciparum*-iRBCs. (A)** Plasma titration curves to determine the optimal plasma dilution factor for the IgG and IgM assay (indicated with a blue and a green arrow, respectively). A pool of endemic plasma samples with high *P. falciparum* parasitemia (*Pf*-infected hyperimmune endemic) from individuals in a sub-cohort within the *InHost* study were used. For the IgM-assay, positive controls (three acute *Pf*-infected non-endemic plasma from a traveler at D0, D21 and D23) and a negative control (naïve non-endemic plasma, Belgian) were added. **(B)** Schematic of gating strategy measuring the surface reactivity of: asexual (tdTom-) and sexual (tdTom+) iRBCs (Hoechst33342+) to IgG (Hoechst33342+/tdTom- or tdTom+/AF488+) or IgM (Hoechst33342+/ tdTom- or tdTom+/PerCP-Cy5.5.+). Cells are first gated for single cells by forward (FSC) and side scatter (SSC). Infected red blood cells (iRBCs) are separated from the non-infected by the DNA stain Hoechst33342. In the iRBCs gate, cells are further separated by the tdTom marker into asexual (tdTom+) and sexual (tdTom+) iRBCs. An anti-human IgG and IgM secondary antibody is used to measure the reactivity to the surface of asexual (tdTom+) and sexual (tdTom+) iRBCs. **(C)** Gating strategy to verify that the parasite cultures contain only red blood cells, positive for the Glycophorin C (GlyC) red blood cell surface marker**. Abbreviations:** asexual iRBCs, trophozoites; CTL, control; D, day(s); IgG and IgM, immunoglobulin G and M; iRBCs, infected red blood cells; *Pf, P. falciparum*; sexual iRBCs, stage I gametocytes.

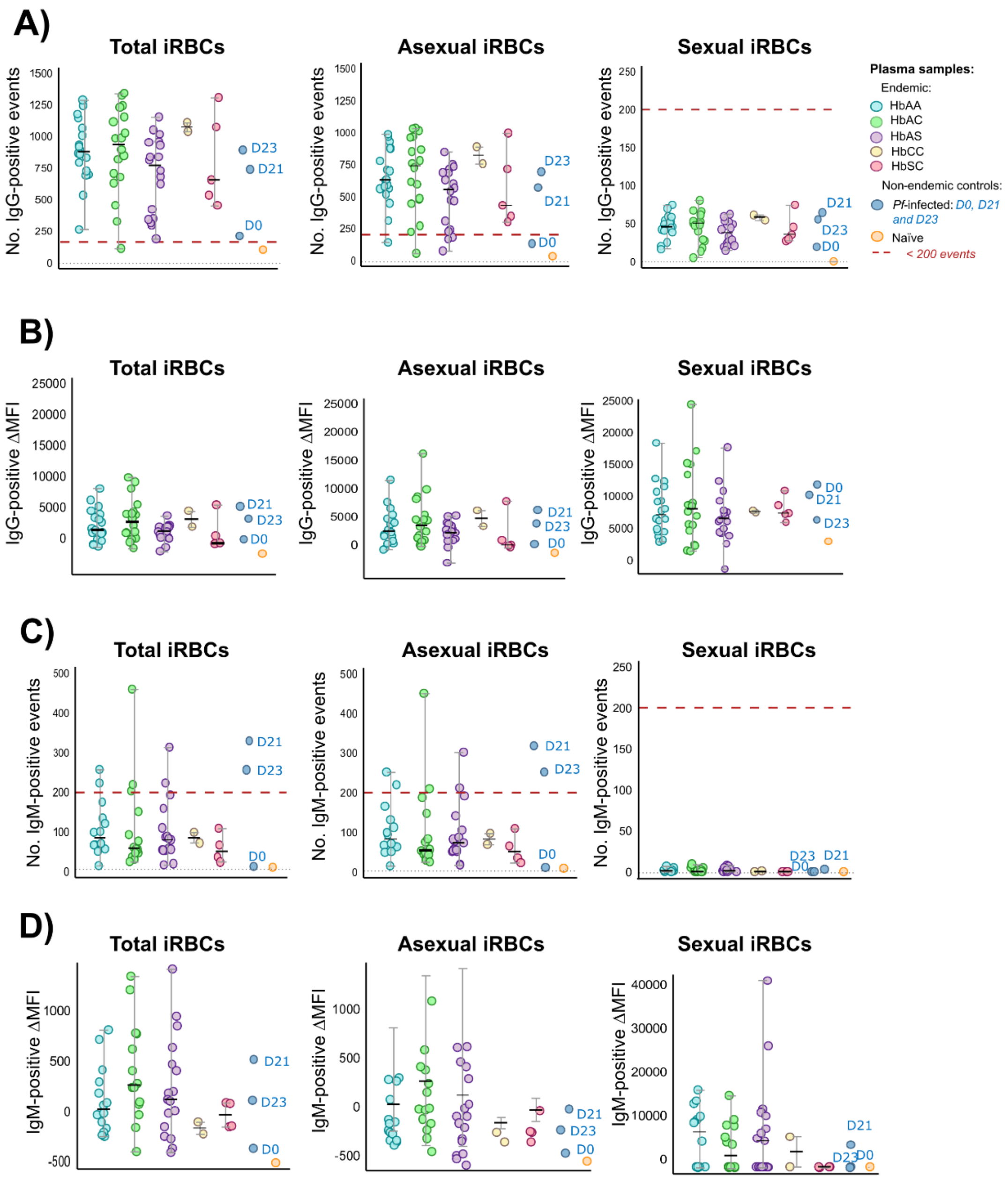


**S5 Fig. IgG- and IgM-mediated responses against *Plasmodium falciparum* trophozoites and stage I gametocytes across HBB genotypes.** Data from *P. falciparum* infected ($n = 44$) and non-infected ($n = 17$) plasma samples collected at the Clinical Research Unit (CRUN) in Nanoro, Burkina Faso (total $n = 61$). **(A)** Event counts and **(B)** MFI of IgG-positive iRBCs: total (left), asexual (middle) and sexual (right) iRBCs. **(C)** Event counts and (**D**) MFI of IgM-positive iRBCs: total (left), asexual (middle) and sexual (right) iRBCs. Positive controls ("*Pf*-infected non-endemic" *i.e.,* three acute *Pf*-infected non-endemic plasma samples from a traveler at D0, D21, and D23) and a negative control (*i.e.,* "Naïve non-endemic plasma" from a volunteer with HbAA genotype) are shown as individual dots per sample. The Y-axis range differs between graphs for visualization. Event counts lower than 200 (red dashed line in **A** and **C)** is not significant and therefore the correspondent MFI plots (in **B** and **D**) are not meaningful data to interpret. No significant differences were found (Kruskal-Wallis with Wilcoxon rank sum post-hoc test for not normally distributed variables, one-way ANOVA with Bonferroni post-hoc for normally distributed variables, $p > 0.05$). **Abbreviations:** asexual iRBCs, trophozoites; Hb, hemoglobin; HBB, human hemoglobin beta; IgG and IgM, immunoglobulin G and M; iRBCs, infected red blood cells; No., number; sexual iRBCs, stage I gametocytes; total iRBCs, trophozoites and stage I gametocytes.

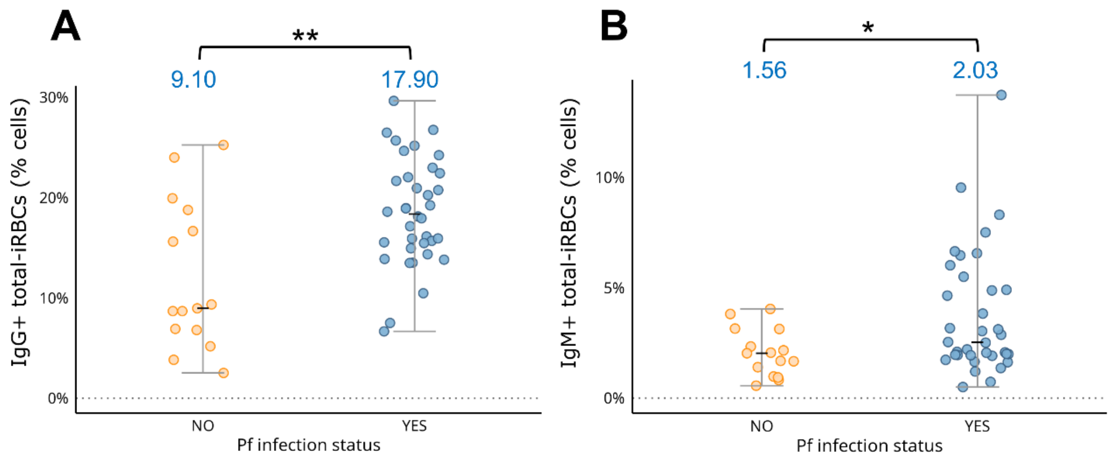


207

**S6 Fig. IgG and IgM positive total iRBCs in plasma from *P. falciparum*-infected and non-infected individuals.** Data from *P. falciparum* infected ($n$ = 44) and non-infected ($n$ = 17) plasma samples collected at the Clinical Research Unit (CRUN) in Nanoro, Burkina Faso (total $n$ = 61). Data points show the median proportions of IgG-positive **(A)** or IgM-positive **(B)** total iRBCs for each "*Pf*-infection" group (infected versus non-infected). The values for each group are indicated in blue above the boxes. The Y-axis range differs between graphs for visualization. Statistical tests used are the Beta-binomial GLMs with Bonferroni adjustment for pairwise comparisons. Significant p-values are shown as: $p < 0.05$*, $p < 0.01$**, $p < 0.001$***; non-significant p-values as: ns. **Abbreviations:** asexual iRBCs, trophozoites; GLMs, generalized linear models; IgG and IgM, immunoglobulin G and M; iRBCs, infected red blood cells; *Pf, P. falciparum*; sexual iRBCs, stage I gametocytes; total iRBCs, trophozoites and stage I gametocytes.

208

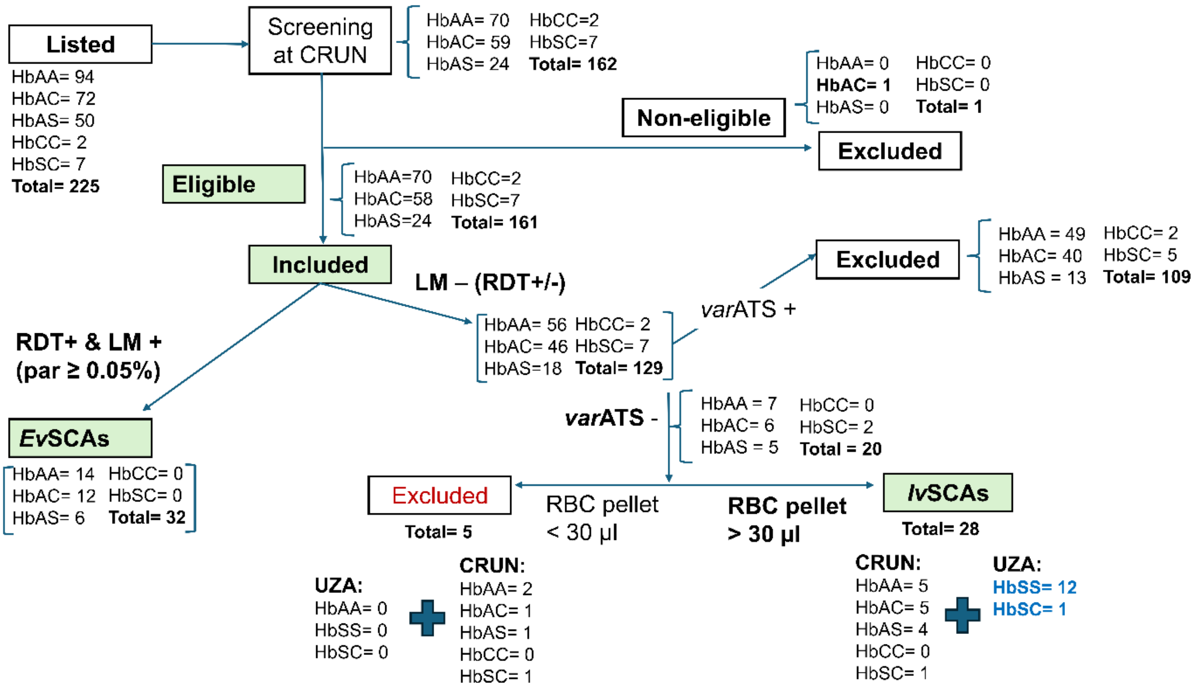


**S7 Fig. Flow chart of participant selection in the *ex vivo* and *in vitro* SC assays.** Participants from the *InHost* study were listed based on their HBB genotype, then screened at CRUN for eligibility and inclusion into this sub-cohort. *P. falciparum* diagnosis was done with RDT, LM and *var*ATS. *P. falciparum* infected samples identified by RDT and LM with a parasitemia ≥ 0.05% were included in the *ex vivo* sexual conversion assays (*ev*SCAs). Non-infected samples (LM- [RDT+ or -]) were confirmed to be true negative with *var*ATS qPCR and were cryopreserved to be used in the *in vitro* sexual conversion assays (*iv*SCAs). Samples with a RBC pellet (recovered after thawing) > 30 µl after dilution with RPMI to 50% Hct were included in the *iv*SCAs. **Abbreviations:** Hb, hemoglobin; HBB, human hemoglobin beta; Hct, hematocrit; LM, light microscopy; *P. falciparum, Plasmodium falciparum;* RBC, red blood cell; RDT, rapid diagnostic test; *var*ATS qPCR, *var* gene acidic terminal sequence (*var*ATS) quantitative PCR (qPCR).

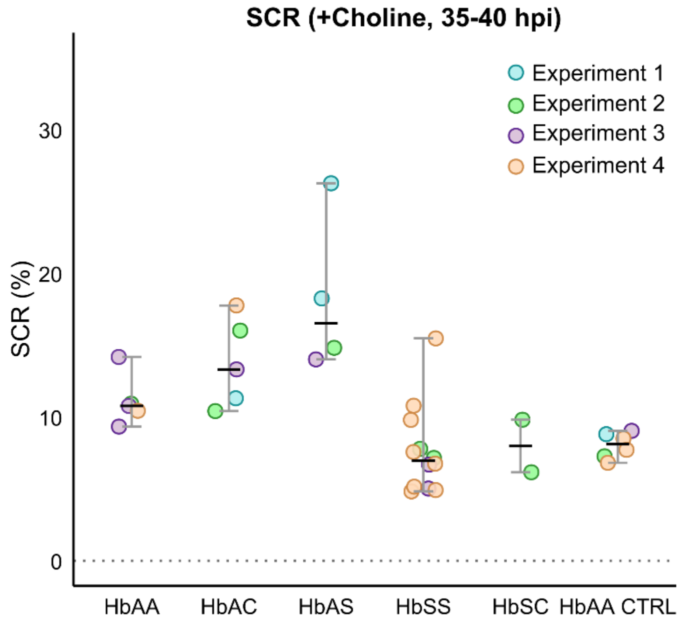


**S8 Fig. Experimental variability of SCR measurements in +Choline cultures, grouped by HBB genotype.** Timepoint of measurement: at 35-40 hpi (Generation 1). **Abbreviations:** CTRL, control sample; Hb, hemoglobin; HBB, human hemoglobin beta.

**S1A Table.** **Statistical analysis outcomes from GLMs or GLMMs and Kruskal-Wallis tests in *ex vivo* SC assays.**

| **Model definition** | **Model Type** | **Estimates (transformed to probability scale / estimated probability** | **Contrasts** | **OR [95% CI] (Wald Z tests)** | **(Adjusted) *p*-value** | **Bayesian OR [credible interval]** | **Comments** |
|---|---|---|---|---|---|---|---|
| **SCR+Choline ~ HBB genotype + (1\|Sample)**<br><br>***Φ ~ HBB genotype*** | **Beta-binomial GLMM** | AA = 0.027<br>AC = 0.048<br>AS = 0.186 | HbAC vs. HbAA | 1.82 [1.17−2.82] | 0.0032 | - | • Technical replicates modelled as random effect.<br>• Genotype-specific dispersion was modelled.<br>• Bayesian model failed to converge.<br>• Bonferroni correction for pairwise comparisons |
| | | | HbAS vs. HbAA | 8.16 [4.34−15.37] | < 0.0001 | - | |

**Abbreviations:** CI, confidence interval; *ev*SCAs, *ex vivo* sexual conversion assays; GLMs or GLMMs, generalized linear (mixed) models; Hb, hemoglobin; HBB, human hemoglobin beta; iRBCs, infected red blood cells; OR, odds ratio; SC, sexual conversion; SCR, sexual conversion rate; Sexual conversion rates (SCRs) are the proportion of *Plasmodium falciparum* parasites that commit to sexual development relative to total parasitemia.

**S1B Table. Statistical analysis outcomes from GLMs or GLMM and Kruskal-Wallis test in the *in vitro* SC assays.**

| Model definition | Model Type | Median[IQR] | Estimates (transformed to probability scale / estimated probability | Contrasts | OR [95% CI] (Wals Z tests) | (Adjusted) *p*-value | Bayesian OR [credible interval] | Comments |
|---|---|---|---|---|---|---|---|---|
| **SCR+Choline (Gen 1, 35-40hpi) ~ HBB genotype** | **Beta-binomial GLM** | - | AA = 0.114<br>AC = 0.138<br>AS = 0.182<br>SS = 0.076<br>SC = 0.081 | HbAC vs. HbAA | 1.25 [0.81−1.92] | 0.77 | 1.25 [0.80−1.85] | • Bonferroni correction for pairwise comparisons |
| | | | | HbAS vs. HbAA | 1.73 [1.12−2.66] | 0.0056 | 1.73 [1.09−2.52] | |
| | | | | HbSS vs. HbAA | 0.64 [0.43−0.95] | 0.021 | 0.65 [0.43−0.92] | |
| | | | | HbSC vs. HbAA | 0.68 [0.35−1.33] | 0.62 | 0.67 [0.28–1.15] | |
| **SCR+Choline (Gen 1, 25-30hpi) ~ HBB genotype** | **Kruskal-Wallis test** | AA= 7.58[0.98)<br>AC= 9.41[5.14]<br>AS= 15.00[6.48]<br>SS= 8.44 [3.76]<br>SC= 7.18 [1.81] | - | - | - | 0.094 | - | • Beta-binomial model assumptions violated (quantile deviations detected). |
| | | | | - | - | | - | |
| | | | | - | - | | - | |
| | | | | - | - | | - | |
| **SCR-Choline (Gen 1, 35-40hpi) ~ HBB genotype** | **Beta-binomial GLM** | - | AA = 0.321<br>AC = 0.326<br>AS = 0.333<br>SS = 0.162<br>SC = 0.124 | HbAC vs. HbAA | 1.02 [0.66−1.59] | 1 | 1.02 [0.65-1.48] | • Bonferroni correction for pairwise comparisons |
| | | | | HbAS vs. HbAA | 1.06 [0.66−1.69] | 1 | 1.06 [0.65-1.56] | |
| | | | | HbSS vs. HbAA | 0.41 [0.27−0.61] | < 0.0001 | 0.41 [0.27-0.58] | |
| | | | | HbSC vs. HbAA | 0.30 [0.14−0.64] | 0.0003 | 0.29 [0.10-0.52] | |
| **SCR-Choline (Gen 1, 25-30hpi) ~ HBB genotype** | **Beta-binomial GLM** | - | AA = 0.250<br>AC = 0.255<br>AS = 0.299<br>SS = 0.149<br>SC = 0.106 | HbAC vs. HbAA | 1.03 [0.65−1.63] | 1 | 1.04 [0.65-1.55] | • Bonferroni correction for pairwise comparisons |
| | | | | HbAS vs. HbAA | 1.28 [0.81−2.03] | 0.73 | 1.29 [0.81-1.93] | |
| | | | | HbSS vs. HbAA | 0.52 [0.35−0.79] | 0.0003 | 0.54 [0.35-0.76] | |
| | | | | HbSC vs. HbAA | 0.36 [0.17−0.77] | 0.0032 | 0.35 [0.13-0.65] | |
| **Late stages proportion (Gen 0, 35-40hpi) ~ HBB genotype** | **Kruskal-Wallis test** | AA= 45.00[20.00]<br>AC= 58.00[10.00]<br>AS=26.00[18.20]<br>SS=9.00[16.50]<br>SC= 8.00[6.00] | - | - | - | 0.05 | - | • Beta-binomial model assumptions violated (dispersion and quantile deviations).<br>• *Not measured for some samples due to limited RBCs; SCR measured for all samples.* |
| | | | | - | - | | - | |
| | | | | - | - | | - | |
| | | | | - | - | | - | |
| **Late stages proportion (Gen 1, 25-30hpi) ~ HBB genotype** | **Kruskal-Wallis test** | AA= 2.00[3.00]<br>AC= 4.00[1.00]<br>AS=3.50[3.50]<br>SS= 2.50[3.25]<br>SC=4.00[1.00] | - | - | - | 0.13 | - | • Beta-binomial model assumptions violated (quantile deviations). |
| | | | | - | - | | - | |
| | | | | - | - | | - | |
| | | | | - | - | | - | |
| **Late stages proportion (Gen 1, 35-** | **Beta-binomial GLM** | - | AA = -1.45<br>AC = -0.21<br>AS = -0.52 | HbAC vs. HbAA | 0.81 [0.24−2.76] | 1 | 0.81 [0.17-2.03] | • Bonferroni correction for pairwise comparisons |
| | | | | HbAS vs. HbAA | 0.60 [0.15−2.31] | 1 | 0.59 [0.09-1.61] | |
| | | | | HbSS vs. HbAA | 0.21 [0.06−0.66] | 0.0026 | 0.22 [0.06-0.55] | |

| | | | | | | | | |
|---|---|---|---|---|---|---|---|---|
| **40hpi) ~ HBB genotype** | | | SS = -1.57<br>SC = -0.63 | HbSC vs. HbAA | 0.53 [0.09−3.02] | 1 | 0.49 [0.01-1.64] | |
| **Parasitemia (Gen 0, 0-5hpi) ~HBB genotype** | **Beta-binomial GLM** | - | AA = -4.99<br>AC = 0.16<br>AS = 0.026<br>SS = 0.11<br>SC = 0.024 | HbAC vs. HbAA | 1.18 [0.40−3.48] | 1 | 1.17 [0.29-2.72] | • Bonferroni correction for pairwise comparisons |
| | | | | HbAS vs. HbAA | 1.03 [0.31−3.37] | 1 | 1.01 [0.19-2.49] | |
| | | | | HbSS vs. HbAA | 1.12 [0.45−2.82] | 1 | 1.15 [0.42-2.49] | |
| | | | | HbSC vs. HbAA | 1.02 [0.24−4.41] | 1 | 0.93 [0.04-2.60] | |
| **Reticulocytes ~HBB genotype** | **Beta-binomial GLM** | - | AA = -4.13<br>AC = -0.14<br>AS = 0.11<br>SS = 0.94<br>SC = 0.85 | HbAC vs. HbAA | 0.86 [0.32−2.36] | 1 | 0.87 [0.27−1.94] | • Bonferroni correction for pairwise comparisons |
| | | | | HbAS vs. HbAA | 1.13 [0.41−3.07] | 1 | 1.11[0.28−2.44] | |
| | | | | HbSS vs. HbAA | 2.56 [1.20−5.44] | 0.008 | 2.54 [1.11−4.87] | |
| | | | | HbSC vs. HbAA | 2.33 [0.82−6.64] | 0.18 | 2.19 [0.44−4.92] | |

**Abbreviations:** CI, confidence interval; GLMs or GLMMs, Generalized Linear (Mixed) Models; Hb, hemoglobin; HBB, human hemoglobin beta; iRBCs, infected red blood cells; IQR, Interquartile range; *iv*SCAs, *in vitro* sexual conversion assays; OR, Odds Ratio; SC, sexual conversion; SCR, sexual conversion rate; Sexual conversion rates (SCRs) are the proportion of *Plasmodium falciparum* parasites that commit to sexual development on the total parasitemia.

**S1C Table .** **Statistical analysis outcomes from GLMs or GLMMs and Kruskal-Wallis tests in the IgG/IgM detection assays.**

| Model definition | Model Type | Estimates | Contrasts | OR [95% CI] (Wald Z tests) | (Adjusted) *p*-value$ | Bayesian OR [credible interval] | Comments |
|---|---|---|---|---|---|---|---|
| **IgG+ total iRBCs (%cells) ~ HBB genotype** | **Beta-binomial GLM** | AA = -1.52<br>AC = -0.10<br>AS = -0.23<br>CC = 0.25<br>SC = -0.031 | HbAC vs. HbAA | 0.90 [0.57−1.43] | 1 | 0.90 [0.59−1.30]<br>0.78 [0.50−1.10]<br>1.23 [0.42−2.27]<br>0.96 [0.47−1.60] | • Bonferroni correction for pairwise comparisons |
| | | | HbAS vs. HbAA | 0.78 [0.50−1.22] | 0.67 | | |
| | | | HbCC vs. HbAA | 1.28 [0.54−3.01] | 1 | | |
| | | | HbSC vs. HbAA | 0.97 [0.49−1.92] | 1 | | |
| **IgG+ asexual iRBCs (%cells) ~ HBB genotype** | **Beta-binomial GLM** | AA = -1.28<br>AC = -0.091<br>AS = -0.30<br>CC = 0.29<br>SC = -0.10 | HbAC vs. HbAA | 0.91 [0.53−1.56] | 1 | 0.92 [0.55−1.37]<br>0.74 [0.46−1.10]<br>1.30 [0.34−2.71]<br>0.89 [0.37−1.59] | • Bonferroni correction for pairwise comparisons |
| | | | HbAS vs. HbAA | 0.74 [0.44−1.25] | 0.61 | | |
| | | | HbCC vs. HbAA | 1.34 [0.49−3.68] | 1 | | |
| | | | HbSC vs. HbAA | 0.91 [0.40−2.04] | 1 | | |
| **IgG+ sexual iRBCs (%cells) ~ HBB genotype** | **Beta-binomial GLM** | AA = -3.70<br>AC = -0.11<br>AS = -0.15<br>CC = 0.27<br>SC = 0.030 | HbAC vs. HbAA | 0.89 [0.59−1.36] | 1 | 0.89 [0.59−1.24]<br>0.86 [0.60−1.20]<br>1.25 [0.49−2.14]<br>1.01 [0.52−1.59] | • Bonferroni correction for pairwise comparisons |
| | | | HbAS vs. HbAA | 0.86 [0.58−1.29] | 1 | | |
| | | | HbCC vs. HbAA | 1.31 [0.63−2.73] | 1 | | |
| | | | HbSC vs. HbAA | 1.03 [0.56−1.89] | 1 | | |
| **IgM+ total iRBCs (%cells) ~ HBB genotype** | **Beta-binomial GLM** | AA = -3.33<br>AC = -0.067<br>AS = -0.039<br>CC = -0.067<br>SC = -0.44 | HbAC vs. HbAA | 0.94 [0.52−1.68] | 1 | 0.94 [0.52−1.44] | • Bonferroni correction for pairwise comparisons |
| | | | HbAS vs. HbAA | 0.96 [0.55−1.68] | 1 | 0.97 [0.58−1.48] | |
| | | | HbCC vs. HbAA | 0.94 [0.29−3.03] | 1 | 0.85 [0.13−1.85] | |
| | | | HbSC vs. HbAA | 065 [0.24−1.72] | 1 | 0.62 [0.17−1.21] | |
| **IgM+ asexual iRBCs (%cells) ~ HBB genotype** | **Beta-binomial GLM** | AA = -3.35<br>AC = -0.076<br>AS = -0.039<br>CC = -0.069<br>SC = -0.42 | HbAC vs. HbAA | 0.93 [0.52−1.66] | 1 | 0.93 [0.54−1.45] | • Bonferroni correction for pairwise comparisons |
| | | | HbAS vs. HbAA | 0.96 [0.55−1.67] | 1 | 0.97 [0.58−1.46] | |
| | | | HbCC vs. HbAA | 0.93 [0.29−3.01] | 1 | 0.84 [1.10−1.79] | |
| | | | HbSC vs. HbAA | 0.65 [0.25−1.72] | 1 | 0.64 [0.17−1.20] | |
| **IgM+ sexual iRBCs (%cells) ~ HBB genotype** | **Beta-binomial GLM** | AA = -7.01e+00<br>AC = 1.41e-01<br>AS = 2.06e-02<br>CC = -4.40e-01<br>SC = -1.97e+01 | HbAC vs. HbAA | 1.15 [0.34−4.00] | 1 | 1.04 [0.28−2.47] | • Bonferroni correction for pairwise comparisons |
| | | | HbAS vs. HbAA | 1.02 [0.32−3.00] | 1 | 0.98 [0.26−2.25] | |
| | | | HbCC vs. HbAA | 0.64 [0.05−9.00] | 1 | 0.52 [0.001−2.44] | |
| | | | HbSC vs. HbAA | 0.00 [0.00−Inf] | 1 | 0.00 [0.00−0.027] | |

| | | | | | | | |
|---|---|---|---|---|---|---|---|
| **IgG+ total iRBCs (%cells) ~ *Pf*-infection** | **Beta-binomial GLM** | NO = -2.06<br>YES = 0.59 | YES vs. NO | 1.81 [1.35−2.43] | **0.001** | 1.82 [1.31−2.43] | • Bonferroni correction for pairwise comparisons<br>• |
| **IgM+ total iRBCs (%cells) ~ *Pf*_infection** | **Beta-binomial GLM** | NO = -3.70<br>YES = 0.41 | YES vs. NO | 1.51 [1.01−2.23] | **0.043** | 1.51 [0.96−2.20] | • Bonferroni correction for pairwise comparisons |

**Abbreviations:** Asexual iRBCs, trophozoite-iRBCs; CI, confidence interval; GLMs or GLMMs, Generalized Linear (Mixed) Models; Hb, hemoglobin; HBB, human hemoglobin beta; iRBCs, infected red blood cells; OR, Odds Ratio; *Pf, P. falciparum*; Sexual iRBCs, stage I gametocyte-iRBCs; Total iRBCs, trophozoite- and stage I gametocyte-iRBCs. %cells are the proportion of IgG- or IgM-positive infected red blood cells.

**S2 Table. . Risk analysis of SCRs and demographic/clinical variables across HBB genotypes in *ev*SCAs.**

| | HbAA | HbAC | HbAS | *p*-value[$] |
|---|---|---|---|---|
| **SCR *n*(%)** | 14 (100) | 12 (100) | 6 (100) | < 0.001 |
| **Gender *n*(%)** | | | | 0.734 |
| Female | 8 (57.1) | 5 (41.7) | 3 (50) | |
| Male | 6 (42.9) | 7 (58.3) | 3 (50) | |
| **Village *n*(%)** | | | | 0.710 |
| Nanoro | 2 (14.3) | 2 (16.7) | 2 (33.3) | |
| Nazoanga | 5 (35.7) | 3 (25) | 1 (16.7) | |
| Soum | 7 (50) | 6 (50) | 2 (33.3) | |
| Séguédin | 0 (0) | 1 (8.3) | 1 (16.7) | |
| **Ethnicity *n*(%)** | | | | 0.231 |
| Gourounsi | 1 (7.1) | 4 (33.3) | 1 (16.7) | |
| Mossi | 13 (92.9) | 8 (66.6) | 5 (83.3) | |
| **Symptomatic *n*(%)** | | | | 0.351 |
| NO | 5 (35.7) | 4 (33.3) | 4 (66.7) | |
| YES | 9 (62.3) | 8 (66.6) | 2 (33.3) | |
| **RDT Positives *n*(%)** | 14 (100) | 12 (100) | 6 (100) | NA |
| **LM Positives *n*(%)** | 14 (100) | 12 (100) | 6 (100) | NA |
| ***P. falciparum* infections *n*(%)** | 14 (100) | 12 (100) | 6 (100) | NA |
| **Mixed Infections = None *n*(%)** | 14 (100) | 12 (100) | 6 (100) | NA |
| **Age, mean (SD)** | 9.31 (7.80) | 7.58 (1.60) | 8.03 (1.66) | 0.957 |
| **Axillary Temperature, mean (SD)** | 36.54 (0.50) | 36.53 (0.43) | 36.37 (0.29) | 0.747 |
| **LM (par/µl), mean (SD)** | 10603.14 (12153.08) | 9653.67 (5609.73) | 16620.33 (21368.16) | 0.820 |
| **LM (gam/µl), mean (SD)** | 0 (0) | 19.58 (48.74) | 13.00 (31.84) | 0.288 |
| ***VarATS* (par/µl), mean (SD)** | 15354.50 (17269.06) | 10262.25 (9008.48) | 36290.50 (47430.62) | 0.750 |

[$]Categorical variables test: Chi Square /Fisher exact test. Continuous variables test: Kruskal Wallis test.
Data from *P. falciparum*-infected samples (parasitemia $\geq$ 0.05% or 2500 parasites/µl) collected in Nanoro, Burkina Faso. Frequencies within each category group are expressed as percentages (%). **Abbreviations:** *ev*SCAs, *ex vivo* sexual conversion assays; Hb, hemoglobin; HBB, human hemoglobin beta; IQR, Interquartile range; LM, Light microscopy; Par, Parasites; RDT, Rapid Diagnostic test; SD, Standard Deviation; SCRs, Sexual conversion rates.

**S3 Table. Median SCRs (IQR) by HBB genotype measured in *iv*SCAs with flow cytometry.**

| | SCRs (%), median (IQR) | | | |
|---|---|---|---|---|
| | +Choline | | -Choline | |
| **HBB genotype** | **25-30 hpi** | **35-40 hpi** | **25-30 hpi** | **35-40 hpi** |
| **HbAA CTRL** | 7.78 (2.59) | 8.12 (1.35) | 22.70 (1.71) | 33.10 (3.15) |
| **HbAA** | 7.58 (0.98) | 10.80 (0.49) | 24.00 (5.36) | 35.50 (7.25) |
| **HbAC** | 9.41 (5.14) | 13.30 (4.71) | 28.60 (7.80) | 33.10 (10.9) |
| **HbAS** | 15.00 (6.48) | 16.50 (5.64) | 29.10 (4.95) | 31.70 (2.32) |
| **HbSS** | 8.44 (3.76) | 6.96 (3.17) | 16.00 (7.53) | 18.40 (5.88) |
| **HbSC** | 7.18 (1.81) | 7.99 (1.83) | 10.20 (1.83) | 11.90 (2.81) |

Timepoints measured: at 25-30 hpi and 35-40 hpi (Generation 1). **Abbreviations:** CTRL, control; Hb, hemoglobin; HBB, human hemoglobin beta; hpi, hours post-invasion; IQR, inter-quartile range; *iv*SCAs, *in vitro* sexual conversion assays.

**S4 Table. Risk analysis of factors across HBB genotype groups in *iv*SCAs.**

| | HbAA | HbAC | HbAS | HbSS | HbSC | *p*-value$ |
|---|---|---|---|---|---|---|
| ***n*(%)** | 5 (100) | 5 (100) | 4 (100) | 12 (100) | 2 (100) | NA |
| **Blood group *n*(%)** | | | | | | 0.06 |
| O+ | 1 (20) | 2 (40) | 2 (50) | 7 (58) | 1 (50) | |
| A+ | 0 (0) | 0 (0) | 0 (0) | 4 (33) | 0 (0) | |
| B+ | 0 (0) | 2 (40) | 1 (25) | 0 (0) | 0 (0) | |
| A- | 2 (40) | 0 (0) | 0 (0) | 1 (9) | 0 (0) | |
| B- | 2 (40) | 1 (20) | 1 (25) | 0 (0) | 1 (50) | |
| **Reticulocytes, median (IQR)** | 1.42 (0.29) | 1.21 (0.73) | 1.72 (0.65) | 4.55 (2.66) | 3.78 (1.93) | <0.001 |
| **Age, median (IQR)** | 8 (19) | 5 (1) | 48.50 (17.50) | 27 (7.75) | 17.50 (12.50) | <0.05 |
| **Gender *n*(%)** | | | | | | 0.05 |
| Female | 4 (80) | 3 (60) | 4 (100) | 3 (25) | 1 (50) | |
| Male | 1 (20) | 2 (40) | 0 (0) | 9 (75) | 1 (50) | |
| **Village *n*(%)** | | | | | | <0.001 |
| Nanoro | 5(100) | 4(80) | 3(75) | 0 (0) | 1(50) | |
| Nazoanga | 0 (0) | 1(20) | 1(25) | 0 (0) | 0 (0) | |
| Soum | 0 (0) | 0 (0) | 0 (0) | 0 (0) | 0 (0) | |
| Séguédin | 0 (0) | 0 (0) | 0 (0) | 0 (0) | 0 (0) | |
| Other° | 0 (0) | 0 (0) | 0 (0) | 12(100) | 1(50) | |
| **Ethnicity *n*(%)** | | | | | | <0.001 |
| Mossi | 5(45) | 4(80) | 4(100) | 0 (0) | 1 (50) | |
| Gourounsi | 0(0) | 1(20) | 0 (0) | 0 (0) | 0(0) | |
| Other° | 0 (0) | 0 (0) | 0 (0) | 12 (100) | 1 (50) | |

$Categorical variables test: Fisher's exact test, Continuous variables test: Kruskal Wallis test. °Other country of origin than Burkina Faso or other ethnicity outside Burkina Faso (not specified) *i.e.,* samples from the UZA. Data from *P. falciparum* non-infected samples collected in Nanoro, Burkina Faso and the UZA (Belgium). Frequencies within each category group are expressed as percentages (%). **Abbreviations:** Hb, hemoglobin; HBB, human hemoglobin beta; hpi, hours post-invasion; IQR, Interquartile range; *iv*SCAs, *in vitro* sexual conversion assays ; SCR, Sexual conversion rates; UZA, Antwerp University Hospital.

**S5 Table. RBC counts per µl analyzed during flow cytometry to exclude hemolysis of fragile mutant RBCs.**

| HBB genotype | RBCs/µl (Median[IQR]) | | |
|---|---|---|---|
| | **35-40 hpi (Gen0)** | **25-30 hpi (Gen1)** | **35-40 hpi (Gen1)** |
| **HbAA** | 9901[8,884] | 10,309[753] | 9,009[11,102] |
| **HbAC** | 13,699[998] | 14,493[12,425] | 17,857[19,491] |
| **HbAS** | 17,857[2,660] | 16,621[3,622] | 17,908[6,241] |
| **HbSS** | 13,246[3,619] | 13,524[6,229] | 16,270[10,272] |
| **HbSC** | 14,925[0] | 10,736[4,889] | 17,801[5,454] |
| ***p*** | 0.44 (ns) | 0.67 (ns) | 0.57 (ns) |

Measurements were done at three timepoints: 35-40 hpi (Generation 0), and 25-30 hpi and 35-40 hpi (Generation1), for HbAC; HbAS, HbSS and HbSC RBCs. Statistical test: Kruskal-Wallis. Non-significant (ns) p-values are indicated in the table ($p > 0.05$). **Abbreviations:** Gen, generation; Hb, hemoglobin; HBB, human hemoglobin beta; hpi, hours post-invasion; RBCs, red blood cells.

**S6 Table. Analysis of parasite stage development by individual parasite culture within each HBB genotype group.**

| Sample No | Sample type | HBB Genotype | PMR | Late stages 35-40 hpi (Generation 0) (%) | Late stages 25-30 hpi (Generation 1) (%) | Late stages 35-40 hpi (Generation 1) (%) | Fold change 1* | Fold change 2 £ | PARASITE DEVELOPMENT PATTERN | SCR (%) 35-40 hpi (Generation 1) |
|---|---|---|---|---|---|---|---|---|---|---|
| 1 | HbAA CTRL | HbAA | 3 | 60.00 | 10.05 | 33.57 | 0.2 | 3.3 | EXPECTED DEVELOPMENT | 8.81 |
| 2 | HbAA CTRL | HbAA | 3 | 46.74 | 7.69 | 25.27 | 0.2 | 3.3 | EXPECTED DEVELOPMENT | 7.27 |
| 3 | HbAA CTRL | HbAA | 5 | 11.32 | 2.02 | 19.81 | 0.2 | 9.8 | EXPECTED DEVELOPMENT | 9.04 |
| 4 | HbAA CTRL | HbAA | 4 | 50.00 | 5.09 | 12.98 | 0.1 | 2.6 | EXPECTED DEVELOPMENT | 8.51 |
| 5 | HbAA CTRL | HbAA | 5 | 42.86 | 5.54 | 14.36 | 0.1 | 2.6 | EXPECTED DEVELOPMENT | 7.72 |
| 6 | HbAA CTRL | HbAA | 4 | 40.91 | 5.07 | 12.70 | 0.1 | 2.5 | EXPECTED DEVELOPMENT | 6.82 |
| 7 | HbAA | HbAA | 3 | 52.81 | 6.79 | 34.10 | 0.1 | 5.0 | EXPECTED DEVELOPMENT | 10.93 |
| 8 | HbAA | HbAA | 3 | 21.99 | 1.68 | 35.68 | 0.1 | 21.2 | EXPECTED DEVELOPMENT | 10.78 |
| 9 | HbAA | HbAA | 1 | 45.16 | 1.66 | 18.73 | 0.04 | 11.3 | EXPECTED DEVELOPMENT | 9.34 |
| 10 | HbAA | HbAA | 2 | 70.41 | 5.29 | 24.41 | 0.1 | 4.6 | EXPECTED DEVELOPMENT | 10.44 |
| 11 | HbAA | HbAA | 5 | 8.22 | 0.00 | 20.92 | 0.0 | NA | EXPECTED DEVELOPMENT | 14.19 |
| 12 | HbAC | HbAC | 2 | 61.66 | 12.44 | 28.33 | 0.2 | 2.3 | DELAYED DEVELOPMENT | 11.31 |
| 13 | HbAC | HbAC | 3 | 51.43 | 4.29 | 31.83 | 0.1 | 7.4 | EXPECTED DEVELOPMENT | 10.42 |
| 14 | HbAC | HbAC | 1 | 60.67 | 3.79 | 8.72 | 0.1 | 2.3 | DELAYED DEVELOPMENT | 16.02 |
| 15 | HbAC | HbAC | 3 | 7.08 | 2.45 | 13.95 | 0.3 | 5.7 | DELAYED DEVELOPMENT | 13.32 |
| 16 | HbAC | HbAC | 2 | 57.69 | 4.58 | 27.49 | 0.1 | 6.0 | EXPECTED DEVELOPMENT | 17.77 |
| 17 | HbAS | HbAS | 3 | 40.16 | 5.22 | 14.52 | 0.1 | 2.8 | EXPECTED DEVELOPMENT | 26.27 |
| 18 | HbAS | HbAS | 3 | 32.86 | 5.91 | 15.90 | 0.2 | 2.7 | EXPECTED DEVELOPMENT | 18.26 |
| 19 | HbAS | HbAS | 2 | 18.67 | 1.70 | 5.14 | 0.1 | 3.0 | DELAYED DEVELOPMENT | 14.82 |
| 20 | HbAS | HbAS | 2 | 8.94 | 1.43 | 3.27 | 0.2 | 2.3 | (PARTIAL) ARREST/EXTREMELY DELAYED DEVELOPMENT | 14.02 |
| 21 | HbSS | HbSS | 2 | 28.81 | 5.60 | 16.41 | 0.2 | 2.9 | DELAYED DEVELOPMENT | 7.79 |
| 22 | HbSS | HbSS | 2 | 20.59 | 4.76 | 15.26 | 0.2 | 3.2 | DELAYED DEVELOPMENT | 7.16 |
| 23 | HbSS | HbSS | 3 | 7.58 | 1.87 | 11.46 | 0.2 | 6.1 | DELAYED DEVELOPMENT | 5.03 |
| 24 | HbSS | HbSS | 4 | 9.84 | 1.81 | 10.66 | 0.2 | 5.9 | DELAYED DEVELOPMENT | 6.69 |
| 25 | HbSS | HbSS | 4 | 4.71 | 3.33 | 1.99 | 0.7 | 0.6 | (PARTIAL) ARREST/EXTREMELY DELAYED DEVELOPMENT | 6.76 |
| 26 | HbSS | HbSS | 2 | 2.70 | 1.61 | 2.13 | 0.6 | 1.3 | (PARTIAL) ARREST/EXTREMELY DELAYED DEVELOPMENT | 4.92 |
| 27 | HbSS | HbSS | 1 | 1.54 | 2.13 | 3.57 | 1.4 | 1.7 | (PARTIAL) ARREST/EXTREMELY DELAYED DEVELOPMENT | 15.48 |
| 28 | HbSS | HbSS | 2 | 2.86 | 1.65 | 1.28 | 0.6 | 0.8 | (PARTIAL) ARREST/EXTREMELY DELAYED DEVELOPMENT | 4.83 |
| 29 | HbSS | HbSS | 2 | 20.97 | 6.06 | 9.41 | 0.3 | 1.6 | (PARTIAL) ARREST/EXTREMELY DELAYED DEVELOPMENT | 10.79 |
| 30 | HbSS | HbSS | 2 | 3.16 | 1.33 | 0.76 | 0.4 | 0.6 | (PARTIAL) ARREST/EXTREMELY DELAYED DEVELOPMENT | 5.16 |
| 31 | HbSS | HbSS | 3 | 14.97 | 4.28 | 5.71 | 0.3 | 1.3 | (PARTIAL) ARREST/EXTREMELY DELAYED DEVELOPMENT | 9.81 |
| 32 | HbSS | HbSS | 3 | 19.42 | 8.05 | 11.14 | 0.4 | 1.4 | (PARTIAL) ARREST/EXTREMELY DELAYED DEVELOPMENT | 7.57 |
| 33 | HbSC | HbSC | 1 | 1.77 | 4.76 | 4.67 | 2.7 | 1.0 | (PARTIAL) ARREST/EXTREMELY DELAYED DEVELOPMENT | 9.82 |
| 34 | HbSC | HbSC | 2 | 13.59 | 2.70 | 9.25 | 0.2 | 3.4 | DELAYED DEVELOPMENT | 6.16 |

Sample size *n* per group: HbAA *n* = 5, HbAC *n* = 5, HbAS *n* = 4, HbSS *n* = 12 and HbSC *n* = 2); and RBC samples of donors from a malaria non-endemic area (“HbAA CTRL” *n* = 6). For each parameter (PMR, late stages percentage at 35-40 hpi at Generation 0 and 25-30/35-40 hpi, at Generation 1, and fold change 1 and 2), the median (Q2), interquartile range (Q2-Q3), and full percentile range (Q1-Q4) were calculated in HbAA cultures, which served as the reference. Values from cultures in HbAC, HbAS, HbSS and HbSC genotypes were compared against these distributions and color-coded as follows: green = median value of sample is within the HbAA interquartile range (Q2-Q3) values. An expected decrease median value of sample is within the HbAA Q1 values and an expected increase median value of sample is within the HbAA Q4 values; orange = for an expected decrease median value of sample is within the HbAA Q4 values, and for an expected increase median value of sample is within the HbAA Q1 values, and red = for an expected decrease median value of sample is outside the HbAA Q4 values; for an expected increase median value of sample is outside the HbAA Q1 values. Values in red are outside the expected range (*i.e.,* HbAA full percentile range [Q1-Q4]). Cultures were then classified into three developmental classes according to the number of parameters outside the HbAA reference range: (1) *expected development* (≤ 1 parameter outside), (2) *delayed development* (2-3 parameters outside), and (3) (*partially) arrested/extremely delayed development* (≥ 4 parameters outside). Fold-change 1 was defined as the ratio of late-stage proportions between 35-40 hpi (Generation 0) and 25-30 hpi (Generation 1), with an expected decrease (< 1), whereas fold-change 2 was defined as the ratio between 25-30 hpi and 35-40 hpi (both Generation 1), with an expected increase (> 1). These expected patterns reflect the normal developmental progression in parasites grown in HbAA RBCs. Sexual conversion rates (SCR) at 35-40 hpi (Generation 1) are also shown. **Abbreviations:** CTRL, control sample; Fold change 1, late stages proportion at 25–30 hpi (Generation 1) divided by late stages proportion at 35–40 hpi (Generation 0); Fold change 2, late stages proportion at 35–40 hpi (Generation 1) divided by late stages proportion at 25–30 hpi (Generation 1); Hb, hemoglobin; HBB, human hemoglobin beta; hpi, hours post-invasion; PMR, parasite multiplication rate; SCR, sexual conversion rate.

**S7 Table. Median IgG- and IgM-positive proportions of total, asexual and sexual iRBCs by HBB genotype.**

| | IgG-positive % cells | | | IgM-positive % cells | | |
|---|---|---|---|---|---|---|
| **HBB genotype** | **Total iRBCs** | **Asexual iRBCs** | **Sexual iRBCs** | **Total iRBCs** | **Asexual iRBCs** | **Sexual iRBCs** |
| **HbAA** | 16.80 (5.91) | 21.40 (9.34) | 2.37 (0.66) | 2.17 (1.78) | 3.39 (2.87) | 0.06 (0.12) |
| **HbAC** | 19.10 (8.00) | 25.00 (11.60) | 2.58 (1.40) | 1.50 (2.38) | 2.34 (3.58) | 0.04 (0.28) |
| **HbAS** | 15.70 (9.89) | 18.30 (13.00) | 2.13 (1.30) | 1.96 (1.29) | 2.96 (2.28) | 0.05 (0.12) |
| **HbCC** | 20.60 (0.87) | 25.70 (2.31) | 2.95 (0.19) | 2.09 (0.33) | 3.26 (0.63) | 0.04 (0.04) |
| **HbSC** | 13.40 (10.10) | 14.80 (9.87) | 1.83 (1.02) | 1.27 (1.24) | 2.00 (1.94) | 0.00 (0.00) |

Measurements done by flow cytometry. **Abbreviations:** asexual iRBCs, trophozoites; Hb, hemoglobin; HBB, human hemoglobin beta; IgG and IgM, immunoglobulin G and M; iRBCs, infected red blood cells; sexual iRBCs, stage I gametocytes; total iRBCs, trophozoites and stage I gametocytes.

**S8 Table. Demographics of participants in IgG/IgM detection assays across HBB genotype groups.**

| | HbAA | HbAC | HbAS | HbSS | HbSC | *p*-value$ |
|---|---|---|---|---|---|---|
| ***Pf* infection *n*(%)** | | | | | | 0.80 |
| NO (*Pf*-) | 4 (22) | 5(28) | 5 (28) | 1 (50) | 2 (40) | |
| YES (*Pf*+) | 14 (78) | 13 (72) | 13 (72) | 1 (50) | 3 (60) | |
| **Parasites per µl (median [IQR])** | 1518 (4244) | 286 (1292) | 496 (1363) | 1916 (1916) | 1235 (6353) | 0.54 |
| **Gametocytes in blood *n*(%)** | | | | | | 0.94 |
| NO | 16 (89) | 15 (83) | 17 (94) | 2(100) | 5 (100) | |
| YES | 2 (11) | 3 (17) | 1 (6) | 0 (0) | 0 (0) | |
| **Age *n*(%)** | | | | | | 0.62 |
| 1 to 4 years | 10 (56) | 9 (50) | 12 (67) | 2 (100) | 4 (80) | |
| 5 to 19 years | 8 (44) | 9 (50) | 6 (33) | 0 (0) | 1 (20) | |
| **Gender *n*(%)** | | | | | | 0.23 |
| Female | 9 (50) | 8 (44) | 12 (67) | 0 (0) | 1 (20) | |
| Male | 9 (50) | 10 (56) | 6 (33) | 2 (100) | 4 (80) | |
| **Village *n*(%)** | | | | | | 0.98 |
| Nanoro | 7 (39) | 7 (39) | 7 (39) | 2 (100) | 2 (40) | |
| Nazoanga | 6 (33) | 6 (33) | 8 (44) | 0 (0) | 1 (20) | |
| Soum | 2 (11) | 2 (11) | 1 (6) | 0 (0) | 1 (20) | |
| Séguédin | 3 (17) | 3 (17) | 2 (11) | 0 (0) | 1 (20) | |
| **Ethnicity *n*(%)** | | | | | | - |
| Mossi | 18 (100) | 18 (100) | 18 (100) | 2 (100) | 5 (100) | |

Significant p-values ($p < 0.05$*, $p < 0.01$**, $p < 0.001$***). $Categorical variables test: Fisher's exact test; Continuous variables test: Kruskal Wallis test.

Data was analyzed in *P. falciparum* infected ($n = 44$) and non-infected ($n = 17$) plasma samples collected at the Clinical Research Unit (CRUN) in Nanoro, Burkina Faso within the context of the *InHost* study. The ethnicity of all study participants was "Mossi". Frequencies within each category group are expressed as percentages (%). **Abbreviations:** Hb, hemoglobin; HBB, human hemoglobin beta; IQR, Interquartile range; *Pf, P. falciparum.*

**S9 Table.** **Key resources and primers/probe sequences.**

| Reagent type (species) or resource | Designation | Source or reference | Identifiers | Additional information |
|---|---|---|---|---|
| **Gene (*P. falciparum*)** | *pfap2-g* | PlasmoDB | PF3D7_1222600 | |
| | *gexp02* | PlasmoDB | PF3D7_1102500 | |
| **Gene [Homo sapiens(human)]** | HBB hemoglobin subunit beta | NLM-NCBI | NCBI_Gene:3043 | HBBseq |
| **Cell line (*P. falciparum*)** | NF54-*gexp02*-Tom | PMID:31601834 | | Maintained in culture with 2 mM choline |
| **Commercial assay or kit** | DNA extraction kit: QIAamp DNA Mini Kit | Qiagen | Cat. No. 51306 | |
| **Chemical compound, drug** | ML10 | PMID:28874661 S. Osborne (LifeArc) and D. Baker (LSHTM) | | cGMP-dependent protein kinase inhibitor |
| | Choline chloride | Sigma-Aldrich | Cat. No. C7527 | |
| **Software, algorithm** | BioEdit 7.2., Biological sequence alignment editor | (Informer Technologies) | RRID: SCR_007361 | HBBseq |
| | FlowLogic™ Flow Cytometry Analysis Software version 8.7. | Inivai Technologies™ | Inivai Technologies ™ | Flow cytometry analysis software |
| | Rstudio 2024.12.1 | © 2025 Posit Software, PBC formerly RStudio, PBC | RRID:SCR_000432 | Statistics analysis software |
| **Primers (and probe for *var*ATS qPCR)** | HBB-fwd (5'->3'): GGGTTGGCCAATCTACTCCC<br><br>HBB-rev (5'->3'): TCAAGCGTCCCATAGACTCAC | | RRID:SCR_003095 | HBBseq |
| | *var*ATS Primer-fw (5'-3'): cccatacacaaccaaytgga<br>*var*ATS Primer-rev (5'-3'): ttcgcacatatctctatgtctatct<br>*var*ATS Probe (5'-3'): 6-FAM-trttccataaatggt-NFQ-MGB | Hofmann N,et al., PLoS Med. 2015. [4] | | *var*ATS qPCR |
| **Antibody** | Pfs16 (mouse, monoclonal) | R.Sauerwein,Teun Bousema, Radboud University | 32F717:B02 | IFA (1:400) |
| | Goat anti-mouse IgG (H+L) Alexa Fluor 488 | Abcam | ab150113 | IFA (1:1000) |
| | Alexa Fluor® 488 AffiniPure™ Mouse Anti-Human IgG | Jackson ImmunoResearch Europe | Code: 209-545-098 | Flow Cytometry (1:640) |
| | PerCP/Cy5.5® Anti-Human IgM antibody (Mouse monoclonal [CH2]) | Abcam | ab201295 | Flow cytometry (1:320) |
| | APC anti-human CD236 (Glycophorin C) Recombinant Antibody | Biolegend® | Catalog# 378204 / 100 tests | |
| **Other** | Mounting medium with DAPI, aqueous Fluoroshield | Abcam | ab104139 | IFA |
| | Hoechst 33342 | BD Pharmingen | Cat. No: 561908 | Flow Cytometry (1:500) |
| | Agglutination tests:<br>Seraclone™ Anti-A (ABO1)<br>Seraclone™ Anti-B (ABO2)<br>Seraclone™ Anti-D (RH1) Blend | BIO-RAD | REF :<br>801320<br>801345<br>802032 | |

**Abbreviations:** Hb, hemoglobin; HBB, human hemoglobin beta; IFA, immunofluorescence assay.

**S10 Table: Overview of experiments and number of cryopreserved RBC samples by HBB genotype.**

| Sample by HBB genotype | Experiments | | | | |
|---|---|---|---|---|---|
| | **1** | **2** | **3** | **4** | **Total *n*** |
| **HbAA (CRUN)** | - | 1 | 3 | 1 | 5 |
| **HbAC (CRUN)** | 1 | 2 | 1 | 1 | 5 |
| **HbAS (CRUN)** | 2 | 1 | 1 | - | 4 |
| **HbSS (UZA)** | - | 2 | 2 | 8 | 12 |
| **HbSC (CRUN/UZA)** | - | 2 | - | - | 2 |
| **Total *n*** | **3** | **8** | **7** | **10** | **28** |

**Abbreviations:** Hb, hemoglobin; HBB, human hemoglobin beta; n, sample size; RBC, red blood cells.

.
